# A Class 1 OLD family nuclease encoded by *Vibrio cholerae* is countered by a vibriophage-encoded direct inhibitor

**DOI:** 10.1101/2025.01.06.631583

**Authors:** Kishen M. Patel, Kimberley D. Seed

## Abstract

Bacteria are constantly threatened by their viral predators (phages), which has resulted in the development of defense systems for bacterial survival. One family of defense systems found widely across bacteria are OLD (for overcoming lysogeny defect) family nucleases. Despite recent discoveries regarding Class 2 and 4 OLD family nucleases and how phages overcome them, Class 1 OLD family nucleases warrant further study as there has only been one anti-phage Class 1 OLD family nuclease described to date. Here, we identify the *Vibrio cholerae-*encoded Class 1 OLD family nuclease Vc OLD and describe its disruption of genome replication of the lytic vibriophage ICP1. Furthermore, we examine its *in vitro* activity, identifying Vc OLD as a DNA nickase. Finally, we identify the first direct inhibitor of a Class 1 OLD family nuclease, the ICP1-encoded Oad1. Our research further illuminates Class 1 OLD family nucleases’ role in phage defense and explores the dynamic arms race between *V. cholerae* and its predatory phage ICP1.

## Introduction

Bacteriophages (phages) are viruses that antagonize their bacterial host to propagate. In natural environments, phages outnumber their bacterial hosts by an estimated ten to one (1). This immense selective pressure has driven the evolution of over 150 distinct bacterial defense systems discovered to date (2, 3). Interestingly, despite their diversity, many bacterial immune systems consist of various combinations of similar protein domains with conserved functions (4). For instance, nuclease domains are seen widely across distinct defense systems, such as ZorE in the Zorya II system and PtuA in the Septu defense system (4, 5). The Septu system also encodes PtuB, an ATPase domain-containing protein similar to those found in other defense systems like the Phage Anti-Restriction-Induced System (PARIS) 1 and PARIS 2 (4, 6, 7). Notably, even within a group or family of defenses, variations in domain composition can give rise to differences in system activity.

OLD (overcoming lysogeny defect) family nucleases exemplify a family of defense systems with variations in domain composition and architecture. A conserved feature of these systems is the presence of both an ABC ATPase domain and a topoisomerase/primase (TOPRIM) domain (8–10). However, the genetic synteny of the genes encoding these domains varies, leading to the classification of OLD family nucleases into four classes. Class 1 OLD family nucleases are single gene systems encoding a protein with an N-terminal ATPase domain and a C-terminal TOPRIM domain. Similarly, Class 2 OLD family nucleases (also known as GajA in the Gabija system) encode a protein with both domains but are associated with a second downstream gene, *gajB*, which encodes a UvrD/ PcrA/Rep-like helicase (11). Class 3 OLD family nucleases function as part of a tripartite system, otherwise known as a retron (8, 9), in which the gene encoding the ATPase-TOPRIM protein is flanked by a reverse transcriptase and non-coding RNA. The most divergent are Class 4 OLD family nucleases encoded by PARIS. In this system, the ATPase (AriA) and TOPRIM (AriB) domains are encoded as separate proteins, though examples exist where the genes are fused, resembling the other classes of OLD family nucleases (8, 9, 12–14).

While recent studies have investigated the function of OLD family nucleases in more complex systems such as Gabija and PARIS, comparatively little is known about Class 1 OLD family nucleases. To date, the only characterized anti-phage Class 1 OLD family nuclease is P2 OLD, encoded by the temperate phage P2, which inhibits the acquisition and induction of the temperate phage lambda in *Escherichia coli* (15). Interestingly, while lambda DNA replication is impeded, degradation of the phage genome was not detected during infection (15). Genetic and biochemical studies of *E. coli* lysogenized by P2 and lambda revealed that P2 OLD reduces the aminoacylation of specific tRNAs and translation efficiency upon lambda induction, ultimately impeding the lambda phage life cycle (16). To investigate how P2 OLD restricts lambda, escape mutants of lambda were generated in *E. coli* in the presence of P2 OLD. These escape phages consistently had mutations in genes that interfere with the host recombination proteins, RecBCD (15, 17, 18). Supporting this, P2 OLD induces cell death in RecBC mutant *E. coli* (19, 20). Thus, the model for P2 OLD is that inhibition of RecBCD leads to P2 OLD activity characterized by cell death and inhibition of the lambda lifecycle (15, 20). Biochemical studies of P2 OLD revealed it has robust nonspecific deoxyribonuclease activity and weak ribonuclease activity *in vitro* (21). Similarly, another Class 1 OLD family nuclease from *Thermus scotoductus* (Ts OLD) exhibits nonspecific deoxyribonuclease activity *in vitro*, though its capacity to inhibit phages has not been assayed (19). Furthermore, *in vitro*, both P2 OLD and Ts OLD have ATPase activity. *In vivo*, P2 OLD has been shown to require its ATPase catalytic residues for activity (19). However, the connection between these *in vitro* and *in vivo* activities is not entirely clear, underscoring the need for the exploration of diverse Class 1 OLD family nucleases.

Here, we identify a *Vibrio cholerae*-encoded Class 1 OLD family nuclease (Vc OLD) that inhibits the unrelated vibriophages ICP1, ICP2, and ICP3. We show that Vc OLD disrupts ICP1 genome replication *in vivo* and, unlike P2 OLD, functions independently of RecBCD. Additionally, we demonstrate that Vc OLD acts as a DNA nickase *in vitro*, with its DNA cleavage activity unaffected by ATP, similar to Ts OLD. Finally, we describe a RecBCD-independent mechanism by which a phage can evade a Class 1 OLD family nuclease, identifying the ICP1-encoded Oad1 (for OLD-antidefense 1) as a direct inhibitor of Vc OLD. Together, our findings expand the understanding of Class 1 OLD family nucleases in phage defense and how phages can counteract them.

## Materials and Methods

### Bacterial and phage growth conditions

All bacteria (listed in Table S1) were grown in Luria-Bertani (LB) broth or on LB agar plates at 37°C with aeration. All *V. cholerae* strains are derivatives of the strain E7946. Antibiotics, when necessary, were supplemented into the broth or agar at the following concentrations for *V. cholerae*: 100 μg/mL streptomycin, 75 μg/ml kanamycin, 100 μg/mL spectinomycin, and 1.25 μg/mL (LB broth) or 2.5 μg/mL chloramphenicol (LB agar). Chloramphenicol was supplemented at 25 μg/mL for *E. coli* strains XL1-Blue and S17. For Rosetta-gami B *E. coli* strains, LB broth and agar were supplemented with 15 μg/ml kanamycin, 34 μg/mL chloramphenicol, and 12.5 μg/mL tetracycline. If the *E. coli* Rosetta-gami B strain carried a pETDUET plasmid, 50 μg/mL carbenicillin was supplemented to the LB and agar. All strains were stocked in 20% glycerol stored at −80°C. *V. cholerae* strains were grown to stationary phase (OD_600_ greater than 1) in liquid LB broth with aeration from colonies on an agar plate. That same day, strains were back-diluted from stationary phase into fresh LB broth to OD_600_ = 0.05 and grown to the OD_600_ specified for each assay below. *E. coli* strains were grown to stationary phase overnight in liquid LB broth with aeration at 37°C.

All phages used in this study are listed in Table S1. High-titer phage stocks were generated as previously described (22). Briefly, confluent lysis plates were made via soft overlay method supplemented with applicable antibiotics on the phages’ respective host strain. Plates were covered with STE Buffer (1M NaCl, 200 mM Tris-HCl, 100 mM EDTA) and rocked overnight at 4°C. The next day, the STE was collected, subjected to a soft spin (10, 000× g for 20 minutes), transferred to a new tube, and then subjected to a hard spin (26, 000× g for 90 minutes). The pellet that remained was nutated in fresh STE with agitation overnight at 4°C, chloroform treated and cleared of chloroform by centrifugation at 5, 000× g for 15 minutes.

### Strain construction and cloning

#### Plasmid construction

All plasmids used in this study are listed in Table S2. Plasmids in this study were generated by Gibson Assembly or by Golden Gate Assembly and introduced into *E. coli* XL1-Blue. Plasmids were electroporated into *E. coli* S17s for mating with *V. cholerae* or *E. coli* Rosetta-gami Bs. Plasmids for the inducible expression of genes of interest in *V. cholerae* are derivatives of pMMB67EH with a theophylline-inducible riboswitch (riboswitch E), as described previously (23). Plasmids for the inducible expression of genes of interest in *E. coli* Rosetta-gami Bs are derivates of pETDuet.

#### V. cholerae construction

*V. cholerae* mutants and chromosomal expression constructs were made by introducing splice by overlap extension (SOE) PCR products by natural transformation as previously described (24). For engineering deletions in *V. cholerae*, the PCR products contained a spectinomycin resistance marker flanked by *frt* recombinase sites added to 1kB up- and downstream regions of homology to the *V. cholerae* chromosome. Spectinomycin-resistant transformants were cured of their resistance, unless otherwise specified, through the mating and induction of a plasmid carrying FLP recombinase, resulting in an in-frame FRT-marked deletion, unless otherwise stated. Strains were then cured of the plasmid. For the chromosomal expression constructs, SOE PCR products combining the expression system construct described in (25), a kanamycin resistance marker, the gene of interest, and arms of homology to the *lacZ* locus were naturally transformed into *V. cholerae* and plated on kanamycin agar plates. All engineered *V. cholerae* strains were confirmed by Sanger or Nanopore amplicon sequencing and colony purified twice before stocking.

#### Phage mutant construction

Phage gene deletions were constructed as previously described (26). In brief, an editing template with the desired deletion was cloned into a vector and introduced to *V. cholerae*. The resulting strain was infected with ICP1, and plaques that formed on this strain were picked into STE Buffer. These ICP1 mutant phages were then used to infect *V. cholerae* with an induced Type 1-E CRISPR-Cas system containing a plasmid with a spacer targeting the phage gene of interest. Recombinant phages were purified three times on this targeting host, and the mutation of interest was confirmed by Sanger sequencing before high-titer stocks were made.

#### Phage spot plates and plaque assays

*V. cholerae* strains were grown in LB broth (supplemented with antibiotics where appropriate) to an OD_600_ = 0.3 or to an OD_600_ = 0.2 and then induced with 1.5mM theophylline and 0.01mM β-D-1-thiogalactopyranoside (IPTG) and/or arabinose at concentrations listed in the figure legends for twenty minutes. For spot plates, *V. cholerae* was added to 0.5% molten LB agar, vortexed, poured on solid agar plates (supplemented with appropriate inducer and antibiotics), and allowed to solidify. Once cooled, 3µL spots of 10-fold serial dilutions of phage were added, allowed to dry, incubated overnight at 37°C, and imaged. For plaque assays, *V. cholerae* was mixed with pre-diluted phage samples. Phage were allowed to attach to *V. cholerae* for seven to ten minutes prior to plating in 0.5% molten top agar (supplemented with antibiotics and inducer where appropriate). Plates were incubated for at least six hours at 37°C. Resulting individual plaques were counted, and the EOP was then calculated by comparing the number of plaques formed on the permissive *V. cholerae* strain relative to the number of plaques formed on a strain of interest. ImageJ was used for plaque size quantification.

#### qPCR for genome replication efficiency

Fold change in phage genome copy number was calculated as previously described (27). In brief, *V. cholerae* were grown to an OD_600_ = 0.2 and then induced with 1.5mM theophylline and 1% (w/v) arabinose and grown to OD_600_ = 0.3 at 37°C with aeration. At an OD_600_ = 0.3, strains were infected with ICP1 at a multiplicity of infection (MOI) of 1, aliquoted across multiple pre-warmed tubes, and returned to 37°C with aeration. A sample was taken and boiled for 10 minutes immediately after adding phage to serve as the starting value. Every four minutes, single tubes were removed from the incubator, 100µL was collected and boiled for 10 minutes, and the remainder 1mL was added to 1mL of ice-cold methanol for deep sequencing (below). Boiled samples were diluted 1:500 to serve as the template for qPCR. All samples were run in biological quadruplicate with technical replicates. Fold change was then calculated as the quantified DNA amount at each timepoint relative to the quantified DNA amount immediately following the addition of phage. Genome replication efficiency was calculated by dividing the fold change in phage genome copy number in a Vc OLD-expressing *V. cholerae* strain by the fold change in phage genome copy number in an empty-expressing control and multiplying by 100.

#### Post-infection deep sequencing and coverage mapping

Once strains were added to ice-cold methanol (above), infected cells were pelleted, washed once with cold 1x phosphate-buffered saline, and stored at −80°C. Pellets were thawe, d and total DNA was extracted using a Qiagen DNeasy blood and tissue DNA purification kit. Briefly, samples were treated as follows: resuspended in 180µL ATL and 40µL proteinase K, incubated for twenty minutes at 56°C, added to 4µL of RNase A, added to 400µL of AL and 400µL of ethanol, bound to the column, washed twice, and eluted in ddH_2_O. SeqCenter (Pittsburgh, PA) performed Illumina sequencing at 200Mb depth. Reads were mapped to reference sequences, and average coverage was calculated across the sequenced biological replicates. The percent of the total mapped reads across each genetic element (*V. cholerae* chromosome I and II and ICP1) was normalized by element length to calculate the mapped reads distribution. DNA-seq reads mapping was done as previously documented (28). Reads were mapped to a reference sequence, and the mean read coverage was calculated from three biological replicates.

#### Growth assays

*V. cholerae* strains were back-diluted to OD_600_ = 0.05 and grown to OD_600_ = 0.3 in LB broth at 37°C with aeration. Strains were then serially diluted, and 5µL were spotted onto agar plates supplemented with inducer (1% (w/v) arabinose and 1.5mM theophylline) and grown overnight at 37°C.

#### Purification of Vc OLD and mass spectrometry

*E. coli* Rosetta-gami Bs containing the pETDuet vector engineered to express Vc OLD-were grown to stationary overnight, back-diluted to OD_600_ = 0.05, grown to OD_600_ = 0.4 to 0.6, and induced with 1mM IPTG for four to six hours in LB broth at 37°C with aeration with appropriate antibiotics. Cells were then pelleted, resuspended in Lysis Buffer A (50mM Tris-HCl pH=7.4, 500mM NaCl, 5% glycerol, 20mM imidazole, protease inhibitor, and 1mM β-mercaptoethanol (BME)) and frozen at −80°C. Cell pellets were then thawed, sonicated, and centrifuged at 12, 000 x g for 30 minutes, transferred to a new conical tube, and centrifuged again. The resulting lysate was loaded onto a column packed with HisPur nickel-nitrilotriacetic acid (Ni-NTA) resin pre-equilibrated with ice-cold Buffer A. After two passes over the column, the flowthrough was discarded, and the column was washed. First, the column was washed with 10 column volumes of Buffer B (50mM Tris-HCl pH=8.0, 1M NaCl, 5% glycerol, 50mM imidazole, and 1mM BME), 2 column volumes of Buffer C (50mM Tris-HCl pH=7.4, 100mM NaCl, 5% glycerol, 150mM imidazole, and 1mM BME), 2 column volumes of Buffer D (50mM Tris-HCl pH=7.4, 100mM NaCl, 5% glycerol, 250mM imidazole, and 1mM BME), 2 column volumes of Buffer E (50mM Tris-HCl pH=7.4, 100mM NaCl, 5% glycerol, 350mM imidazole, and 1mM BME), and eluted in 6 column volumes of Buffer F (50mM Tris-HCl pH=7.4, 100mM NaCl, 5% glycerol, 500mM imidazole, and 1mM BME). Aliquots of the flowthrough of washes and elutes were boiled in Laemmli buffer supplemented with BME and assessed by SDS-PAGE/Coomassie staining. Fractions with a single ∼66kDa band corresponding to Vc OLD-HIS by Coomassie were concentrated using Amicon Centrifugal Filter Units with a 10k molecular weight cut off (MWCO) or by dialysis against a Polyethylene glycol (PEG) 8000 solution using a 3.5k MWCO Slide-A-Lyzer dialysis cassette. The final protein sample was buffer exchanged into Storage Buffer (50mM Tris-HCl pH = 7.4, 300mM NaCl, 5% glycerol, and 1mM BME), and assessed by SDS-PAGE/Coomassie staining and mass spectrometry to verify purity. Protein concentration was determined by absorbance at 280nm using an extinction coefficient of 69900L/(mol*cm). The sample was then aliquoted, snap-frozen in liquid nitrogen, and stored at −80 °C until use. For mass spectrometry, purified Vc OLD was precipitated in trichloroacetic acid at a final concentration of 20% overnight in ice at 4°C. The sample was then centrifuged at 18, 000 x g at 4°C and washed three times in 1mL of 0.01 M HCl and 90% acetone. The resulting pellet was allowed to air dry and then submitted to the Vincent J. Coates Proteomics/Mass Spectrometry laboratory on the University of California Berkeley campus for enzymatic digestion and mass spectrometry analysis.

### Mass spectrometry

Mass spectrometry was performed at the Vincent J. Coates Proteomics/Mass Spectrometry Laboratory (PMSL) at the University of California, Berkeley. Multidimensional protein identification technique (MudPIT) was performed as described (29, 30). Briefly, a 2D nano LC column was packed in a 100-μm inner diameter glass capillary with an integrated pulled emitter tip. The column consisted of 10 cm of ReproSil-Gold C18-3μm resin (Dr. Maisch GmbH)) and 4 cm strong cation exchange resin (Partisphere, Hi Chrom). The column was loaded and conditioned using a pressure bomb. The column was then coupled to an electrospray ionization source mounted on a Thermo-Fisher LTQ XL linear ion trap mass spectrometer. An Agilent 1200 HPLC equipped with a split line so as to deliver a flow rate of 1 µL/min was used for chromatography. Peptides were eluted using a 4-step gradient with 4 buffers. Buffer (A) 5% acetonitrile, 0.02% heptafluorobutyric acid (HFBA), buffer (B) 80% acetonitrile, 0.02% HFBA, buffer (C) 250mM NH4AcOH, 0.02% HFBA, (D) 500mM NH4AcOH, 0.02% HFBA. Step 1: 0-80% (B) in 70 min, step 2: 0-50% (C) in 5 min and 0-45% (B) in 100 min, step 3: 0-100% (C) in 5 min and 0-45% (B) in 100 min, step 4 0-100% (D) in 5 min and 0-45% (B) in 160 min. Collision-induced dissociation (CID) spectra were collected for each m/z. Protein identification, quantification and analysis were done with PEAKS (Bioinformatics Solution Inc) with the following parameters: semi-specific cleavage specificity at the C-terminal site of R and K, allowing for 5 missed cleavages, precursor mass tolerance of 3 Da for low resolution LCMS, and fragment ion mass tolerance of 0.6 Daltons. Methionine oxidation and phosphorylation of serine, threonine and tyrosine was set as variable modifications and Cysteine carbamidomethylation was set as a fixed modification. Peptide hits were filtered using a 1% false discovery rate (FDR). The mass spectrometry proteomics data have been deposited to the ProteomeXchange Consortium via the PRIDE (31) partner repository with the dataset identifier PXD058617 and 10.6019/PXD058617.

### Nuclease Assay

Nuclease assays with Vc OLD were carried out in 50mM Tris-HCl pH = 7.4, 50mM NaCl, 10mM MgCl_2_, 10mM MnCl_2_, 10mM CaCl_2_, 10NiCl_2_, and 10mM ZnCl_2_. Frozen Vc OLD samples were thawed and added to the buffer at varying concentrations. Plasmid DNA substrate, a pUC19 derivative, was added at 100ng per 20µL reaction. Reactions were incubated at 30°C for three hours, and then 1µg/mL of Proteinase K was added to each reaction to digest bound protein and incubated at 37°C for 30 minutes. To generate linear or nicked controls, 100ng of vector was digested in 1X CutSmart Buffer for one hour at 37°C with 10 U of BamHI-HF (NEB) or 10 U of Nb.BtsI (NEB) in 20µL reactions, respectively, and then 1µg/mL of Proteinase K was added to each reaction to digest bound protein and incubated at 37°C for 30 minutes. The entire reaction volume was loaded and run on a 0.8% agarose gel stained with GelRed. ImageJ was used to perform densitometry measurements, quantifying the intensity of linear and nicked vectors compared to the total lane intensity. Reported densitometry measurements are the average of three replicate nuclease assays.

### Western Blot

*V. cholerae* strains were grown as listed above for plaque assay conditions. Prior to mixing with phage*, V. cholerae* was collected via centrifugation and resuspended in ZN Lysis Buffer (50mM Tris-HCl pH = 7.4, 150mM NaCl, 1mM EDTA, protease inhibitor, and 0.5% Triton-X-100). Total protein content was assayed via MicroBCA Assay (Thermo) and then mixed with Laemmli Buffer supplemented with BME. Once in Laemmli Buffer, samples were boiled for ten minutes and loaded into an Any-Kd Mini-PROTEAN TGX Precast gel (Bio-Rad) and run. The amount of total protein loaded into each lane of the blot is listed in the respective figure legend. Then, the gel was transferred to a Mini-size PVDF membrane (Bio-Rad) using a Trans-Blot Turbo system (Bio-Rad). This membrane was blocked overnight in 5% milk in Tris-buffered saline (TBS) with Tween 20 (TBST) at 4°C. For lanes probed using multiple antibodies, membranes were cut between 37kDa and 50kDa standards so that each resulting membrane could be probed with a different antibody. Membranes (cut or whole) were then washed three times in TBST, incubated with a primary α-Flag antibody (GenScript) at a 1:1000 dilution, or incubated with a primary α-HA antibody (GenScript) at a 1:2000 dilution for one hour, washed three times in TBST, incubated with a secondary goat α-rabbit-HRP antibody (Bio-Rad) at a 1:3000 dilution for one hour, washed three times in TBS, developed with Clarity Western ECL substrate (Bio-rad) and imaged using a Chemidoc XRS imaging system (Bio-Rad).

### Co-immunoprecipitation

100mLs of *V. cholerae* culture were grown in LB broth with antibiotics and inducer (1% arabinose, 1.5mM theophylline, and 0.01mM IPTG) at 37°C to an OD_600_ = 0.3 from a back-dilution at OD_600_ = 0.05, pelleted by centrifugation, resuspended to an OD_600_ = 25 in Lysis Buffer B (50mM Tris-HCl pH=7.4, 150mM NaCl, 1mM EDTA, 0.5% Triton-X-100, and protease inhibitor), and frozen at −80°C. Pellets were then thawed, a pinch of lysozyme was added, sonicated, and centrifuged at 18, 000 x g for 15 minutes at 4°C. 40µL of lysate was added to Laemmli Buffer supplemented with BME and boiled for 10 minutes. 850µL of lysate were added to 40µL slurry of M2 anti-FLAG Magnetic Resin (Sigma) and subjected to end-over-end rotation for 3 hours at 4°C. The resin was then washed four times: the first two washes consisted of 1mL washes with ice-cold Wash Buffer 1 (50mM Tris-HCl pH=7.4, 300mM NaCl, 1mM EDTA, and 0.05% Triton-X-100) and the second two washes consisted of 200µL of ice-cold Wash Buffer 2 (50mM Tris-HCl pH=7.4 and 150mM NaCl). During each wash, the resin was subjected to end-over-end rotation for five minutes at 4°C. Finally, bound protein was eluted using ice-cold Elution buffer (50mM Tris-HCl pH=7.4, 150mM NaCl, and 450ng/µL 1x FLAG peptide (GenScript), where 50µL of Elution buffer was added to the resin and subjected to end-over-end rotation for 30 minutes at 4°C. 45µL of the eluates were collected and added to Laemmli Buffer supplemented with BME and boiled for 10 minutes. Samples were then probed for the presence or absence of proteins of interest via Western Blot (above).

### Bioinformatic Analyses

#### Viral Proteomic Tree Generation

Fasta files for phages of interest were uploaded to ViPTree (32), and resulting trees were used in appropriate figures.

#### Gene Graphs

Gene graphs were generated using the Python libraries: DNA features viewer, Biopython, and Matplotlib.

#### Average Nucleotide Identity and Clinker Analysis

Fasta files for phages of interest were uploaded to FastANI (33) and Clinker (34). For average nucleotide identity, the resulting list is shown in Table S5. The resulting Clinker protein comparison matrixes were analyzed. An abbreviated list of these results is shown in Table S3.

#### Predicted Protein Structure

The primary sequence of Vc OLD and Gp205^2018^ were submitted to ColabFold (35) as described in the text. Protein structure comparison was made in PyMOL using the cealign function.

#### Figure Generation and Editing

Figures were generated and edited using GraphPad PRISM and Adobe Illustrator.

## Results

### The Class 1 OLD family nuclease Vc OLD protects *V. cholerae* from diverse phages

Previously, we found that *V. cholerae* isolates from 2008 and 2009 harbored a unique phage defense element (PDE) encoding an OLD family nuclease (protein ID: HAS2513945.1) and the anti-phage toxin-antitoxin system DarTG (Fig.1A) (22). This PDE was necessary and sufficient to completely block plaque formation by the most extensively characterized lytic vibriophage, ICP1 (28, 36, 37). The PDE-encoded OLD family nuclease lacks a downstream helicase or an upstream reverse transcriptase and, therefore, belongs to Class 1 OLD family nucleases (19). Upon deletion of the toxic effector DarT, the efficiency of plaquing (EOP) of the phage ICP1 returned to one, indicating that phage defense by the PDE was largely due to DarTG. Interestingly, we observed that ICP1 plaque size was still reduced, indicating that additional PDE-encoded gene(s) were restricting ICP1 (Fig. 1B). Given that the only other predicted defense encoded by the PDE is Vc OLD, we hypothesized that this phenotype was due to Vc OLD. Thus, we constructed a PDE^+^ Δ*darT* Δ*old* strain and observed plaque size returned to that seen in the permissive, PDE^-^ host, indicating that Vc OLD also contributes to phage defense by the PDE in *V. cholerae* (Fig. 1C).

**Figure 1:**
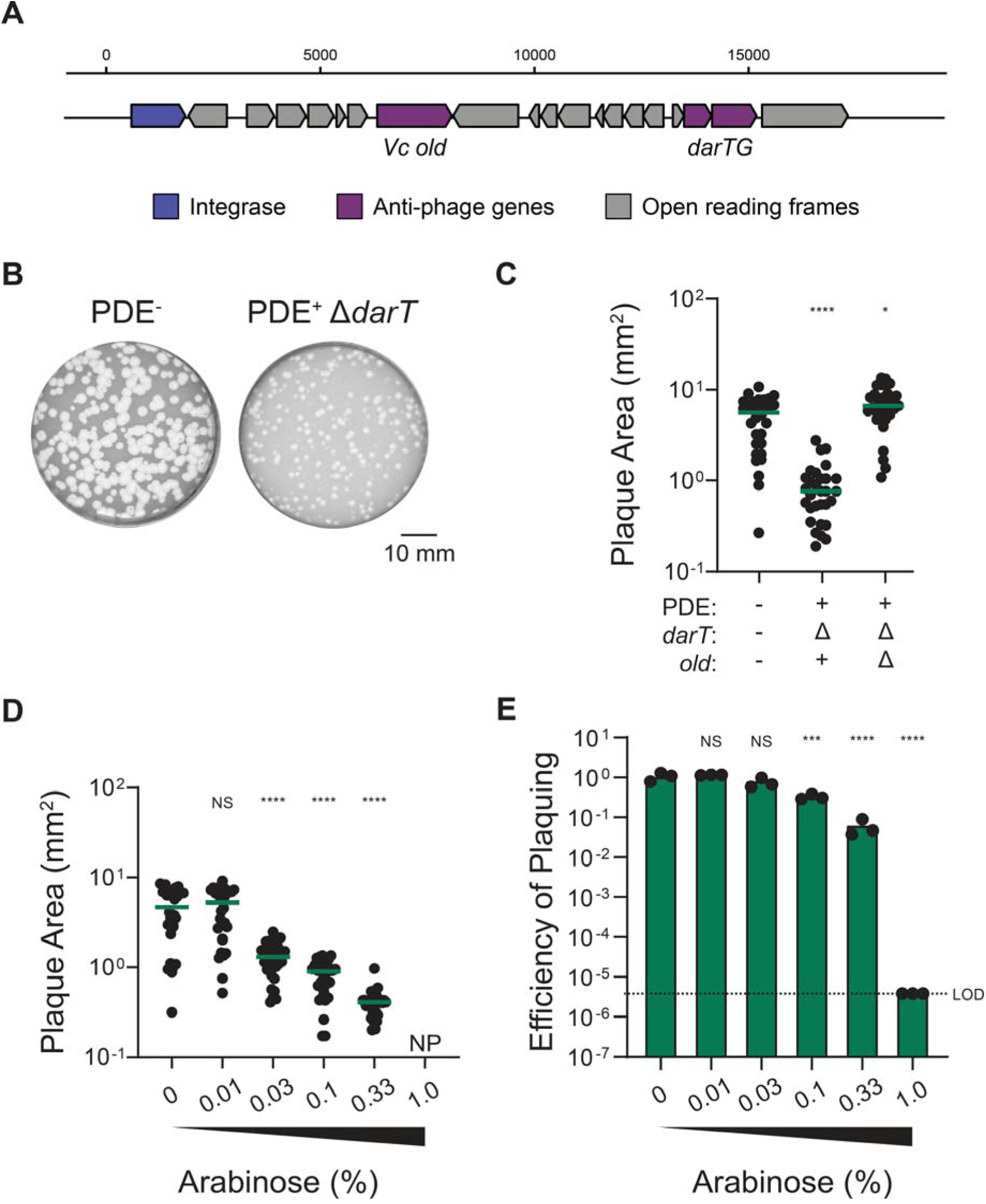
A Class 1 OLD family nuclease (Vc OLD) encoded by a phage defensive element (PDE) in *V. cholerae* restricts the lytic vibriophage ICP1. A) Genetic map (drawn to scale) of the PDE encoded by clinical *V. cholerae* isolates (22). The anti-phage systems *Vc old* and *darTG* are colored in purple, while the predicted *integrase* is shown in deep blue. Other open reading frames not predicted to defend against phages are gray. B) Representative plaque plates (n=3) of ICP1 at the same dilution on otherwise isogenic permissive PDE^-^ and PDE^+^ Δ*darT V. cholerae*. White zones of clearing are ICP1 plaques. The opaque background is the *V. cholerae* lawn. The scale bar is shown in the bottom right. C) The plaque area (mm^2^) of ICP1 plaques on PDE^+/-^ *V. cholerae* and its mutant derivatives. Statistical analyses are the results of analysis of variance (ANOVA) followed by Dunnett’s T3 multiple comparison test using the PDE^-^ *V. cholerae* strain as the control condition (\*\*\*\**P*<0.0001 and \**P*=0.0108). D) The plaque area (mm^2^) of ICP1 on *V. cholerae* expressing Vc OLD at different percentages (w/v) of inducer (arabinose). NP represents no plaques. Statistical analyses are the results of ANOVA followed by Dunnett’s T3 multiple comparison test using the 0% arabinose induction condition as the control condition (\*\*\*\**P*<0.0001, NS = not significant). For B-C, each dot represents an individual plaque. Ten plaques from each biological replicate for three biological replicates (30 plaques total) were measured. The green line indicates the mean of the plaque areas. E) The efficiency of plaquing of ICP1 on *V. cholerae* expressing Vc OLD at different percentages (w/v) of arabinose. Each dot represents an individual biological replicate (n=3). The bar indicates the mean of biological replicates. LOD represents the limit of detection. Statistical analyses are the results of ANOVA followed by Dunnett’s T3 multiple comparison test using the 0% arabinose induction condition as the control condition (\*\*\*\**P*<0.0001, \*\*\**P*=0.0001, NS = not significant).

To determine if Vc OLD is sufficient to limit ICP1 plaquing outside of the PDE, we cloned *Vc old* under an inducible promoter into the chromosome of a permissive PDE^-^ *V. cholerae* strain. Here, we saw that Vc OLD restricted ICP1 in a dose-dependent manner (Fig. 1D). At a high dose of inducer (1% w/v arabinose), Vc OLD completely abolished ICP1 plaquing below our limit of detection (Fig. 1E). Given this robust inhibition, future assays were conducted at this dose of inducer unless otherwise specified. To monitor Vc OLD protein abundance, we introduced a C-terminal HA tag to Vc OLD (herein Vc OLD-HA). Notably, we did not observe any phenotypic differences with Vc OLD after incorporating the HA tag (Fig. S1A and S1B). Then, by western blot, we monitored Vc OLD-HA protein abundance at three induction levels: levels that yielded no ICP1 inhibition, levels that yielded ICP1 inhibition similar to that of Vc OLD in its native context in the PDE, and levels that completely inhibited ICP1. As expected, more induction of Vc OLD-HA led to a greater abundance of protein and more significant inhibition of ICP1 (Fig. S1C).

Prior to this study, P2 OLD was the only characterized Class 1 OLD family nuclease with documented anti-phage activity, and it had been shown to inhibit only a single phage (15). The extent to which Class 1 OLD family nucleases can inhibit unrelated phages, therefore, remains unknown. Previously, we found that ICP2 and ICP3, other phages co-isolated with *V. cholerae* from cholera-positive patient stool samples, were minimally or not inhibited by the PDE, respectively (36). To investigate if Vc OLD could also restrict other phages, we challenged cells expressing Vc OLD at its most restrictive level for ICP1 with ICP2 and ICP3. ICP2 and ICP3 were moderately inhibited by Vc OLD in terms of EOP and plaque size compared to ICP1, indicating that Vc OLD can restrict unrelated phages when overexpressed (Fig. 2). Together, we conclude that Vc OLD is likely lowly expressed in its native locus and is more restrictive to diverse phages ICP1, ICP2, and ICP3 at higher expression levels.

**Figure 2:**
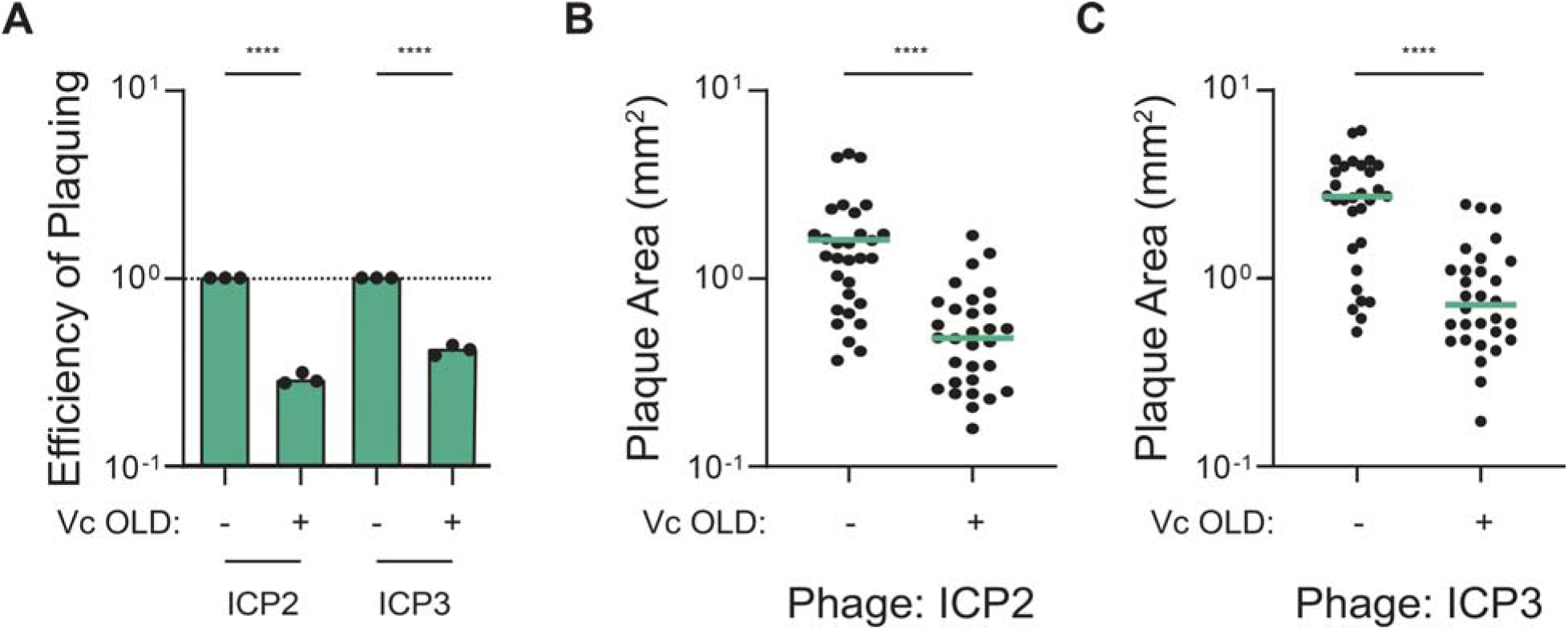
Vc OLD is sufficient to inhibit the unrelated vibriophages ICP2 and ICP3. A) The efficiency of plaquing of ICP2 and ICP3 in *V. cholerae* expressing an empty expression cassette (Vc OLD^-^) or Vc OLD at 1% arabinose. Each dot represents an individual biological replicate (n=3). The bar indicates the mean of biological replicates. An EOP of 1 has been denoted with a dashed line. B) The plaque area (mm^2^) of ICP2 plaques on *V. cholerae* expressing an empty expression cassette (Vc OLD^-^) control or Vc OLD at 1% arabinose. C) The plaque area (mm^2^) of ICP3 in *V. cholerae* expressing a Vc OLD^-^ control or OLD at 1% arabinose. For B-C, each dot represents an individual plaque. Ten plaques from each biological replicate for three biological replicates (30 plaques total) were measured. For A-C, statistical analyses are the results of unpaired *t*-test (\*\*\*\**P*<0.0001).

### Vc OLD inhibits ICP1 genome replication

We hypothesized that Vc OLD was inhibiting ICP1’s genome replication because P2 OLD has been shown to restrict lambda phage prior to genome replication. To test this, we monitored ICP1’s genome replication during infection of *V. cholerae* in the presence and absence of Vc OLD at several time points using qPCR. Previous studies reported that ICP1 completes genome replication within twenty minutes (28, 38). However, we noted that ICP1 genome replication kinetics appeared delayed in the presence of arabinose. Therefore, we extended our analysis to assess replication dynamics over a twenty-four-minute timeframe. By qPCR, we detected a severe reduction in ICP1 genome replication efficiency after sixteen minutes in the presence of Vc OLD (Fig. 3A).

**Figure 3:**
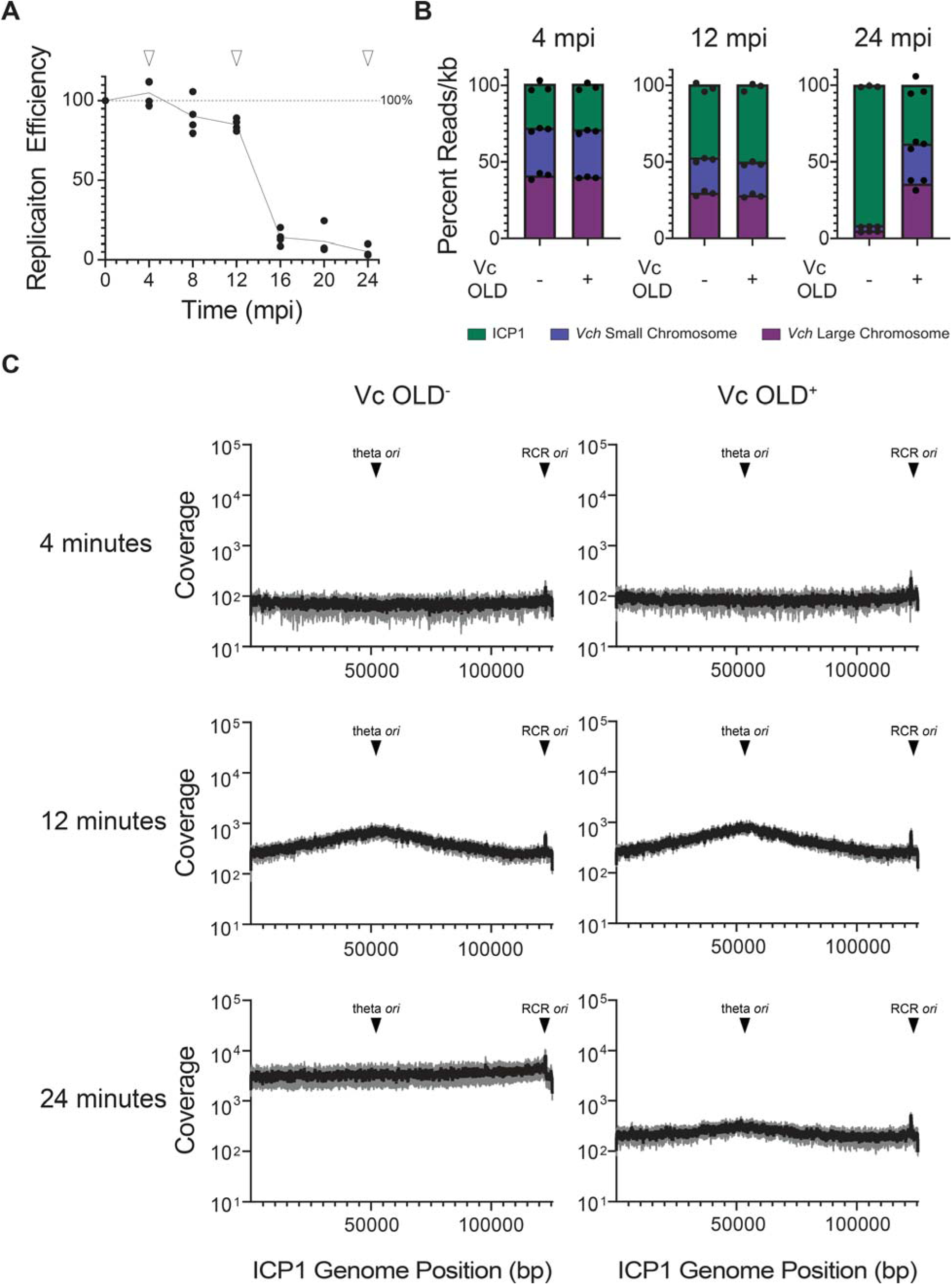
Vc OLD disrupts ICP1 genome replication A) ICP1 genome replication efficiency during infection of *V. cholerae* expressing OLD at 1% arabinose as determined by qPCR. The efficiency of replication is relative to an induced OLD^-^ control. White triangles indicate sample time points selected for deep sequencing. For A-B, each data point is representative of a single biological replicate (n=4). The line connecting each time point represents the average genome replication efficiency over time minutes post-infection (mpi). B) Deep sequencing reads of an ICP1 infection of *V. cholerae* at the time points indicated mapped to the three genetic elements (normalized by length) shown as percent distribution. The bar indicates the mean of biological replicates, which are shown as dots (n=3). Vch stands for *V. cholerae*. C) Reads coverage plots across the ICP1 genome during infection of OLD^-^ (left) and OLD^+^ *V. cholerae* (right). For each time point, the average percent reads coverage is shown as a black line (n=3). The standard deviation appears as gray shading around the line. The theta replication origin of replication (*ori*) and rolling circle replication (RCR) *ori* (28) are labeled.

To gain additional insight into how Vc OLD disrupts ICP1’s genome replication, we deep-sequenced total DNA from phage-infected cells at four-, twelve-, and twenty-four-minutes post-infection. In the absence of arabinose, ICP1 initiates bi-directional (theta) replication by eight minutes post-infection and transitions to rolling circle replication by sixteen minutes, generating the essential genome concatemers that serve as the substrate for packaging into progeny virions (28). Here, mapping sequencing reads to the phage genome confirmed that replication occurred between four- and twelve minutes post-infection in both conditions, +/- Vc OLD (Fig. 3B). Moreover, we observed no apparent differences in ICP1’s initiation of theta replication twelve minutes post-infection as we observed a coverage map that aligned with the expected pattern for theta replication (Fig. 3C).

By twenty-four minutes post-infection, however, phage genome replication patterns diverged markedly depending on the expression of Vc OLD. In the absence of Vc OLD, ICP1 successfully infected *V. cholerae*, evidenced by extensive degradation of the bacterial genome and the majority of sequencing reads mapping to ICP1 (Fig. 3B). The genome coverage map also showed a pronounced skew towards the rolling circle origin of replication, consistent with the expected transition to rolling circle replication (Fig. 3C). In contrast, in the presence of Vc OLD, the percentage of reads mapping to ICP1 failed to increase after twelve minutes and instead decreased (Fig. 3B). Furthermore, no skew towards the rolling circle origin of replication was observed in the presence of Vc OLD (Fig. 3C), indicating that the phage’s transition to rolling circle replication was impeded. Taken together, we conclude that Vc OLD inhibits ICP1’s genome replication after the initiation of theta replication, thus blocking the transition to rolling circle replication and abolishing phage production.

### *In vivo* phage defense activity of Vc OLD is dependent on ATPase and nuclease active site residues but independent of perturbations to RecBCD

A notable feature of P2 OLD function is that perturbations to RecBCD—either through lambda-mediated inhibition or mutation of RecBC—result in P2 OLD-mediated cell death. To investigate whether Vc OLD induces a similar phenotype in a RecBCD-deficient background, we assessed the viability of *V. cholerae* expressing Vc OLD in a strain lacking RecBCD. We engineered a spectinomycin resistance marker in place of the *recBCD* locus in strains harboring the inducible Vc OLD or an empty expression cassette as a control. Expression of Vc OLD in this RecBCD-deficient strain did not result in cell death compared to the control (Fig.4A, Fig. S2), suggesting that Vc OLD does not depend on host RecBCD for activation.

**Figure 4:**
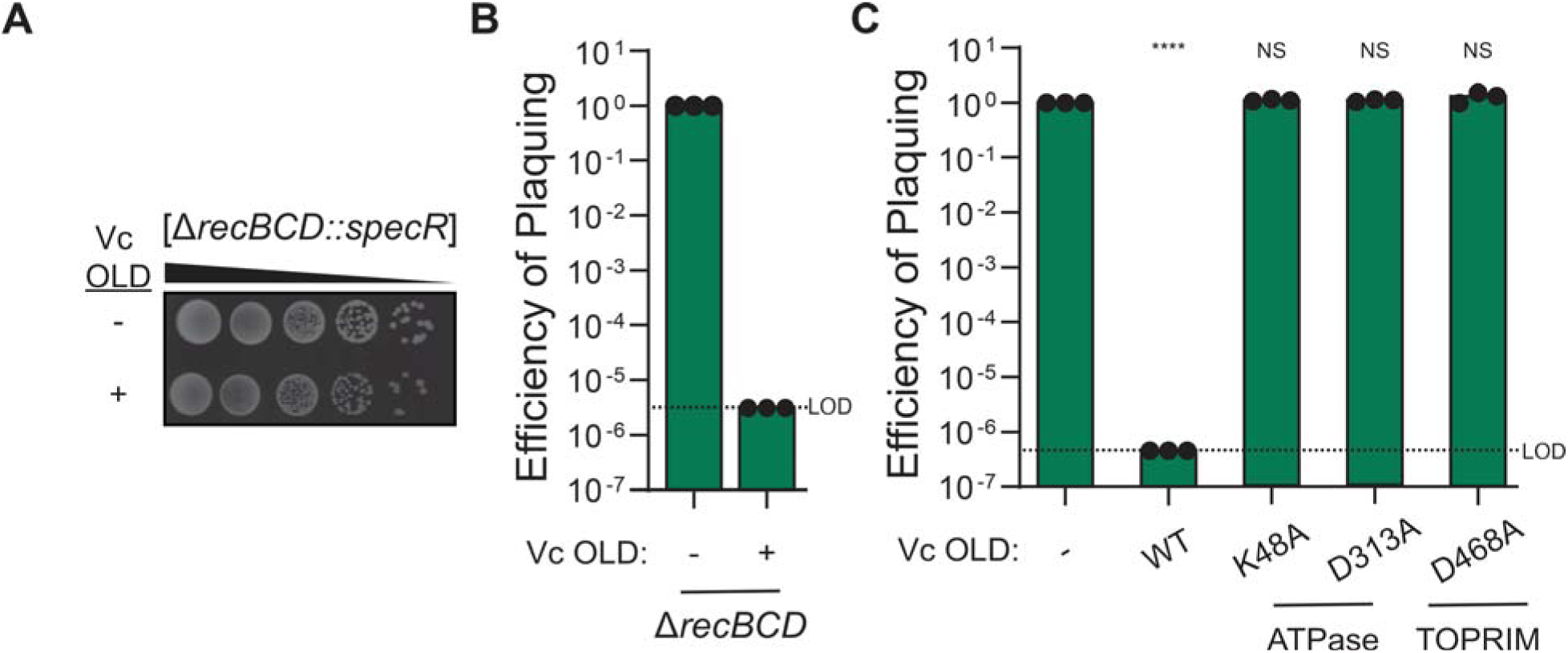
Vc OLD does not require RecBCD in *V. cholerae* and depends on its ATPase and TOPRIM domains for phage defense *in vivo*. A) Representative image of ten-fold serial dilutions of RecBCD^-^ *V. cholerae* expressing or Vc OLD at 1% arabinose or an empty expression cassette (Vc OLD^-^). The black background is the agar plate, while the white spots are colonies. Additional biological replicates are shown in Fig. S2. B) The efficiency of plaquing of ICP1 on RecBCD^-^ *V. cholerae* expressing Vc OLD or a Vc OLD^-^ control induced at 1% arabinose. Statistical significance by unpaired *t*-test is unable to be calculated as the standard deviation is 0.0 at the limit of detection. C) The efficiency of plaquing of ICP1 on *V. cholerae* expressing Vc OLD and Vc OLD mutants at 1% arabinose. WT stands for wildtype. Statistical analyses are the results of ANOVA followed by Dunnett’s T3 multiple comparison test using the Vc OLD^-^ *V. cholerae* strain as the control condition (\*\*\*\**P*<0.0001, NS = not significant). For B-C, each dot represents an individual biological replicate (n=3). The bar indicates the mean of biological replicates. LOD represents the limit of detection.

The current model for P2 OLD-mediated inhibition of lambda phage infection posits that lambda expresses Gam, a RecBCD inhibitor, which subsequently activates P2 OLD and triggers cell death. To test whether Vc OLD can inhibit phages independently of RecBCD, we examined its ability to defend against ICP1 in a RecBCD-deficient background. We found that Vc OLD successfully inhibited ICP1 in the absence of RecBCD (Fig. 4B). These findings indicate that, in contrast to P2 OLD, Vc OLD’s response to phage infection does not rely on RecBCD.

Previously, P2 OLD was shown to require its ATPase and TOPRIM domains to induce cell death in RecBC-deficient *E. coli* (19, 39); however, the requirement of these domains for phage defense was never tested. As such, we next assessed the necessity of the Walker A (required for ATP binding) and Walker B (required for ATP hydrolysis) motifs and/or catalytic residue in the nuclease domain for Vc OLD-mediated phage defense. Because Vc OLD shares low amino acid identity with the previously studied P2 and Ts OLDs (<25%), we identified key residues in these domains by comparing the ColabFold (35) predicted structure of Vc OLD to the 3D crystal structure of Ts OLD (Fig. S3). This comparison enabled us to identify key residues in the Walker A motif (K48),

Walker B motif (D313), and nuclease domain (D468). Using site-directed mutagenesis, we introduced alanine substitutions at these positions to generate three mutant constructs of Vc OLD. We then assessed their ability to inhibit ICP1 through plaque assays. Our results revealed that Vc OLD requires functional Walker A and Walker B motifs as well as the predicted catalytic nuclease residue D468 to inhibit phage *in vivo* (Fig. 4C).

### Vc OLD functions as a nickase *in vitro*

Both P2 OLD and Ts OLD have been shown to have robust but nonspecific DNA cleavage activity *in vitro* (19, 21). Notably, the cleavage activities of both respond differently to ATP: Ts OLD’s activity is unaffected by the addition of ATP, whereas P2 OLD’s DNA cleavage activity is enhanced with the addition of ATP (19, 21). Given our findings that Vc OLD-inhibits phage genome replication (Fig. 3) and requires both ATPase and nuclease domains for phage defense *in vivo* (Fig. 4C), we hypothesized that Vc OLD could cleave DNA. Thus, we sought to directly determine if Vc OLD can cut DNA by purifying Vc OLD with a C-terminal 6xHIS tag and performing nuclease assays with and without the addition of ATP.

To confirm protein purity, we analyzed the purified Vc OLD protein by Coomassie staining (Fig. S4) and mass spectrometry (Table S4), which revealed minimal contaminants and no evidence of contaminating nucleases. We then incubated Vc OLD with circular plasmid DNA to evaluate its nuclease activity, specifically assessing whether it could nick, create double-stranded breaks, or fully degrade the DNA. As controls, we treated the plasmid with BamHI (to linearize the DNA) or Nb.BstI (to generate nicked DNA) to distinguish between these activities. Interestingly, and in contrast to what was found for other Class 1 OLD family nucleases, we observed that VC OLD could nick plasmid DNA but did not induce robust DNA degradation, even at high concentrations of protein (Fig. 5). Moreover, similar to Ts OLD, the addition of ATP did not enhance nicking activity (Fig. S5). These results demonstrate that Vc OLD nicks DNA *in vitro* and that this activity is independent of the addition of ATP.

**Figure 5:**
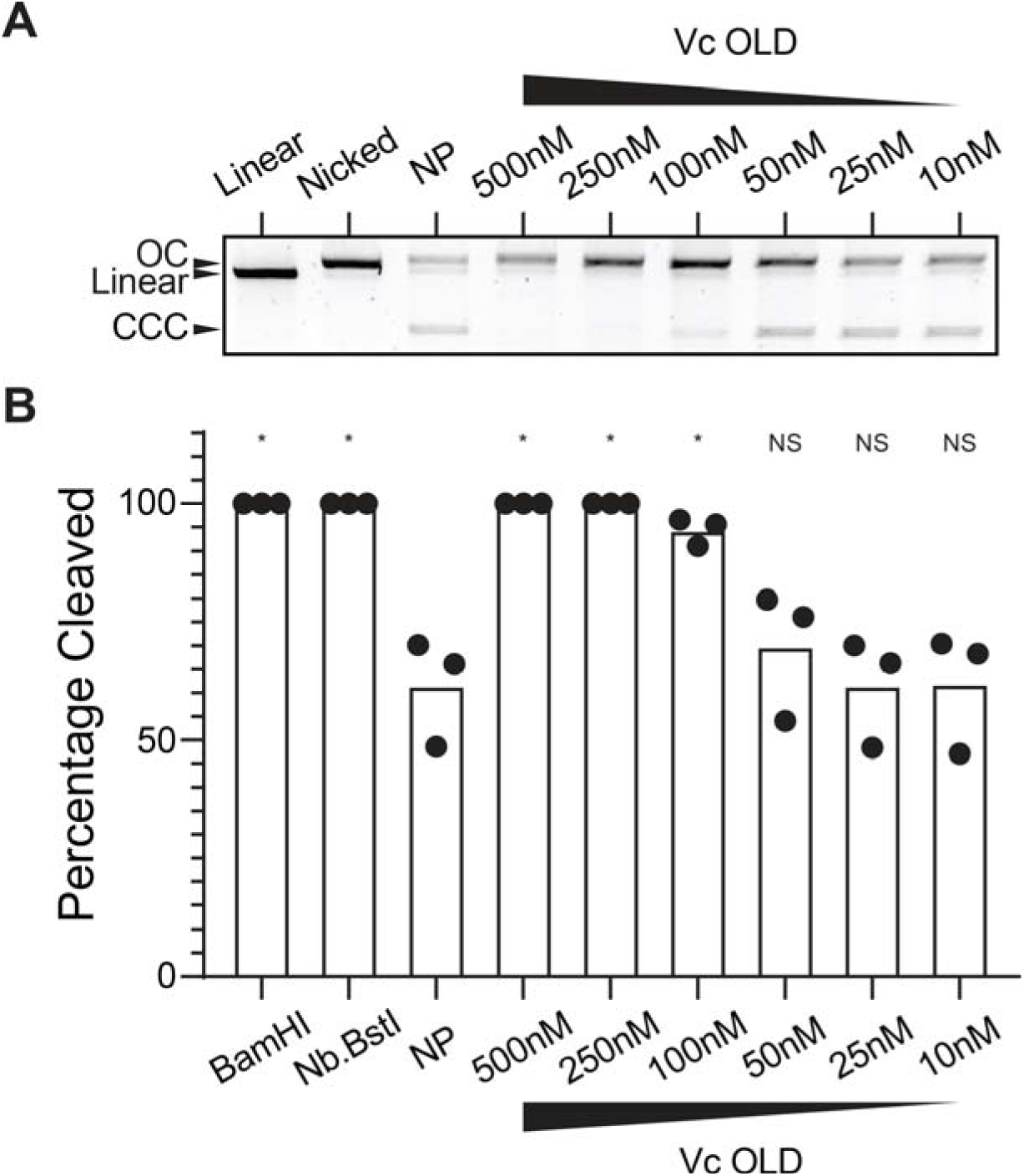
Vc OLD nicks DNA *in vitro*. A) *In vitro* cutting of a plasmid by purified Vc OLD at concentrations indicated above the gel. Plasmid linearized or nicked by restriction enzymes BamHI and Nb.BstI, respectively, are shown in the first two lanes. This image is representative of three replicates, which are quantified in Fig. 5B. NP stands for no protein. B) Densitometry measurements calculating the percent of the plasmid cleaved by control enzymes BamHI and Nb.BstI and by Vc OLD at the different indicated concentrations. Each dot represents an independent replicate (n=3). The bar indicates the mean of replicates. Statistical analyses are the results of ANOVA followed by Dunnett’s T3 multiple comparison test using the no protein (NP) lane as the control condition (\**P*<0.001, NS = not significant).

### ICP1 can encode an OLD anti-defense protein that antagonizes Vc OLD-mediated phage defense

Phages can counter-adapt to defense systems that limit their propagation by mutating the target of the defense system or expressing an inhibitor. In lambda, phages that overcome P2 OLD have mutations in the lambda *red* operon that interferes with *E. coli* recombination machinery, RecBCD (18). Given that Vc OLD activity is RecBCD-independent (Fig. 4), we anticipated ICP1 would escape using a different, uncharacterized mechanism. We first attempted to evolve escape mutants by propagating ICP1_2006, the most well-characterized isolate, on Vc OLD expressed at its most inhibitory concentration. However, this approach failed to yield mutants insensitive to Vc OLD. We then leveraged our collection of 67 genetically distinct isolates of ICP1 (37) to identify any that could naturally overcome Vc OLD. Representatives from the four major clades of ICP1 (ICP1_1992, ICP1_2006, ICP1_2011, and ICP1_2018) (37) were tested on Vc OLD expressing *V. cholerae* (Fig. 6A, Fig. S6). We discovered that ICP1_2018 could plaque robustly on cells expressing Vc OLD, while all other isolates were inhibited.

**Figure 6:**
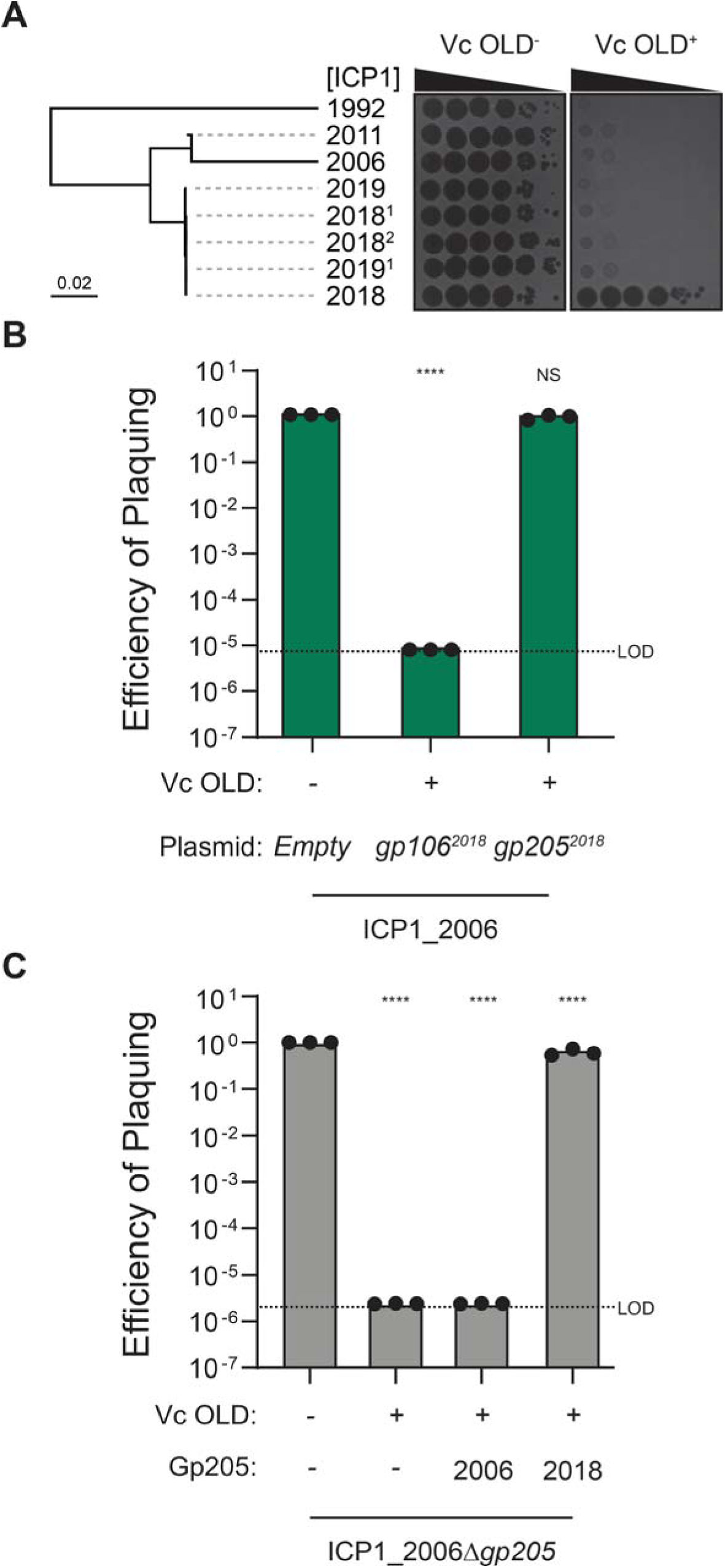
ICP1-encoded Gp205^2018^ allows for escape from Vc OLD. A) Phylogenetic tree of the listed ICP1 isolates (black branches) comparing all translated open reading frames to determine overall genome similarity [based on the tBLASTx algorithm from ViPTree (32)] (left). Representative images (n=3) of ten-fold serial dilutions of the ICP1 isolates indicated spotted on Vc OLD^+/-^ expressing *V. cholerae* strains induced at 1% arabinose (right). Black zones of clearing are plaques. The opaque background is the *V. cholerae* lawn. Other replicates are in Fig. S6. Phage names have been abbreviated for simplicity; the full details of phage strains used are included in Table S1. B) The efficiency of plaquing of ICP1_2006 on Vc OLD^+/-^ expressing *V. cholerae* induced at 1% arabinose and a vector expressing the Gp listed from ICP1_2018. Statistical analyses are results of ANOVA followed by Dunnett’s T3 multiple comparison test using the empty vector containing Vc OLD^-^ *V. cholerae* strain as the control condition (\*\*\*\**P*<0.0001, NS = not significant). C) The efficiency of plaquing of ICP1_2006Δ*gp205* in Vc OLD^+/-^ expressing *V. cholerae* induced at 1% arabinose and an empty vector or a vector expressing Gp205^2006^ or Gp205^2018^. Statistical analyses are results of ANOVA followed by Dunnett’s T3 multiple comparison test using the empty vector containing, Vc OLD^-^ *V. cholerae* strain as the control condition (\*\*\*\**P*<0.0001). For B and C, each dot represents an individual biological replicate (n=3). The bar indicates the mean of biological replicates. LOD represents the limit of detection.

To understand the genetic basis of this escape, we next tested isolates closely related (>99.95% average nucleotide identity) to ICP1_2018 (Table S5). Remarkably, none of these highly related isolates escaped Vc OLD (Fig. 6A). We hypothesized that a difference in a protein-coding region of ICP1_2018 could be responsible for its escape. Thus, we used Clinker (34) to compare the genomes of ICP1_2018 and Vc OLD-sensitive ICP1 isolates, identifying three unique variants of gene products (Gps 106, 205, and 207) encoded by ICP1_2018 (Table S3). We next attempted to clone each of the genes from ICP1_2018 onto a plasmid under an inducible promoter. To this end, we successfully engineered plasmids expressing Gps 106 and 205, but not Gp207. Using the plasmid-expressed genes, we assessed their ability to protect ICP1_2006 from Vc OLD-mediated restriction. Notably, only Gp205 from ICP1_2018 (herein Gp205^2018^) allowed ICP1_2006 to overcome Vc OLD (Fig. 6B).

Gp205 is an uncharacterized protein found in the core genome of all ICP1 isolates (37). Given that an allele of Gp205 is encoded by all isolates of ICP1 sequenced to date, we hypothesized that Gp205 could be essential for ICP1 and a target of Vc OLD. Interestingly, Gp205^2018^ differs from Gp205 in ICP1_2006 (herein Gp205^2006^, protein ID: AXQ70829.1) by only four amino acids, which could account for the escape from Vc OLD. To test the essentiality of Gp205^2006^, we engineered an ICP1_2006 mutant lacking Gp205 (ICP1_2006Δ*gp205*). Surprisingly, this mutant was viable and remained sensitive to Vc OLD, indicating that Gp205^2006^ is not essential or a necessary target of Vc OLD (Fig. 6C). We therefore hypothesized that Gp205^2018^ is a gain-of-function allele providing counter-defense against Vc OLD. To test this, we expressed Gp205^2018^ or Gp205^2006^ alongside Vc OLD and infected strains with ICP1_2006Δ*gp205*. Expression of Gp205^2018^, but not Gp205^2006^, restored the EOP of ICP1_2006Δ*gp205* to approximately one (Fig. 6C), providing additional evidence that Gp205^2018^ is sufficient to provide counter-defense against Vc OLD. Lastly, we tested if Gp205^2018^ could protect the unrelated phage ICP3 against Vc OLD. Consistent with our findings in ICP1, Gp205^2018^ increased ICP3 plaque size in the presence of Vc OLD, whereas Gp205^2006^ had no effect (Fig. S7). Together, these data support the hypothesis that Gp205^2018^ is a gain of function allele sufficient to confer counter-defense against Vc OLD.

### ICP1’s OLD anti-defense protein (Oad1) is a direct inhibitor of Vc OLD

Having identified Gp205^2018^ as a counter-defense against Vc OLD, we sought to investigate its mechanism of action. Many characterized phage counter-defenses function as direct inhibitors of their cognate defense systems (40). Given that Gp205^2018^ confers protection to both ICP1_2006 and ICP3 against Vc OLD, we hypothesized that it acts as a direct inhibitor of Vc OLD. We first asked if Gp205 and Vc OLD interacted *in silico* by modeling the predicted structure of Gp205^2018^ and a modeled Vc OLD dimer using ColabFold (35). This analysis identified regions of interaction between Gp205^2018^ and Vc OLD at or near three of the four amino acid residues that distinguish Gp205^2018B^ from Gp205^2006^: S135, K136, and N144 (Fig. 7A). These findings suggest that these residues might contribute to the interaction with Vc OLD and could underlie the differing abilities of Gp205^2018^ and Gp205^2006^ to counteract Vc OLD defense.

**Figure 7:**
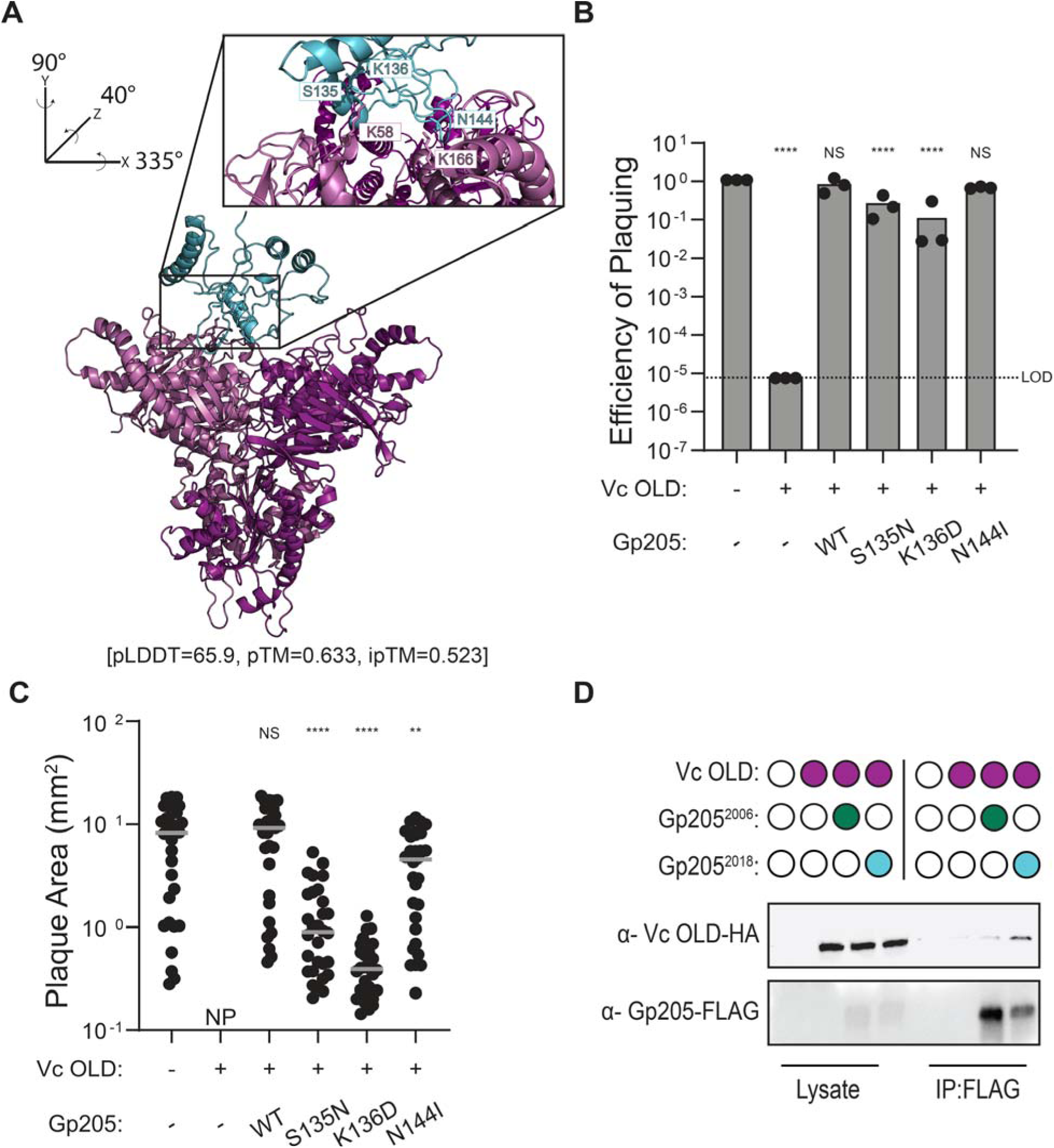
ICP1-encoded Gp205^2018^ (Oad1, for OLD anti-defense 1) binds to Vc OLD. A) The ColabFold (35) predicted structures of Vc OLD modeled as a dimer and Gp205^2018^. Each monomer of Vc OLD is shown in different colors: light pink and purple, while Gp205 is shown in light blue. The pLDDT, pTM, and ipTM scores are written below the structure. The inset was rotated on the axes shown and highlights the predicted interacting residues S135 and N144 from Gp205^2018^ and K58 and K166 from Vc OLD. K136 from Gp205^2018^ is labeled as it is adjacent to S135. These three residues from Gp205^2018^ (S135, K136, and N144) are three of the four amino acids that distinguish Gp205^2018^ from Gp205^2006^. Distances between predicted interacting residues are as follows: 3.0Å between S135 and K58 and 2.1 Å and 3.2Å between N144 and K166 (which has two predicted interactions). B) The efficiency of plaquing of ICP1_2006Δ*gp205* on Vc OLD^+/-^ expressing *V. cholerae* induced at 1% arabinose and an empty vector or a vector expressing Gp205^2018^, or the mutant derivative indicated. Amino acid substitutions revert the Gp205^2018^ residue to the corresponding residue in Gp205^2006^. Each dot represents an individual biological replicate (n=3). The bar indicates the mean of biological replicates. LOD represents the limit of detection. WT stands for wildtype. Statistical analyses are results of ANOVA followed by Dunnett’s T3 multiple comparison test using the empty vector containing Vc OLD^-^ *V. cholerae* strain as the control condition (\*\*\*\**P*<0.0001, NS = not significant). C) The plaque area (mm^2^) of ICP1_2006Δ*gp205* plaques on Vc OLD^+/-^ expressing *V. cholerae* induced at 1% arabinose and an empty vector or a vector expressing Gp205^2018^, or the mutant derivative indicated. Each dot represents an individual plaque. Ten plaques from each biological replicate were measured for three biological replicates (30 plaques total). The grey line indicates the mean of the plaque areas. WT stands for wildtype. Statistical analyses are the results of ANOVA followed by Dunnett’s T3 multiple comparison test using the empty vector containing Vc OLD^-^ *V. cholerae* strain as the control condition (\*\*\*\**P*<0.0001, \*\**P*=0.076, NS = not significant). D) Western blot analysis of the co-immunoprecipitation of Gp205-FLAG from cells expressing the proteins indicated (filled-in circle means being expressed). Lysates used as input for the immunoprecipitation (IP) were probed as controls for expression levels. Images are representatives of two biological replicates; the additional replicate is shown in Fig. S8.

We next evaluated whether the three predicted interacting amino acids that differ between Gp205^2018^ and Gp205^2006^ are necessary for counter-defense against Vc OLD. To this end, we engineered single amino acid reversions in Gp205^2018^ to their corresponding residues in Gp205^2006^ (S135N, K136D, and N144I) and evaluated counter-defense activity against Vc OLD using plaque assays (Fig 7B-C). While none of the individual mutations completely abolished Gp205^2018^’s capacity to confer counter-defense against Vc OLD, the S135N and K136D mutations led to reduced counter-defense, as evidenced by decreased EOP and smaller plaque sizes (Fig. 7B-C). In contrast, the N144I mutant did not alter the EOP but produced smaller plaques (Fig. 7B-C). These findings indicate that while these residues contribute to Gp205^2018^’s counter-defense, no single residue is strictly necessary for its function against Vc OLD.

To experimentally determine if Vc OLD and Gp205 directly interact, we next performed an *in vivo* co-immunoprecipitation assay. We co-expressed C-terminal 3xFLAG tagged Gp205 (Gp205-FLAG) from ICP1_2018 or ICP1_2006 along with VC OLD-HA and incubated the cell lysate with anti-FLAG resin. We observed increased enrichment of Vc OLD-HA with Gp205^2018^-FLAG but not with Gp205^2006^-FLAG (Fig. 7D, Fig. S8). From these data, we conclude that Gp205^2018^ and Vc OLD directly interact. Taken together, these data establish Gp205 as a direct inhibitor of a Class 1-OLD family nuclease, and thus, we propose that Gp205 be named Oad1 (for OLD anti-defense 1).

## Discussion

The flux of mobile genetic elements encoding phage defense systems in *V. cholerae* shapes interactions with phages that prey on this pathogen in the context of disease (41). Notably, in its native locus, Vc OLD provides only modest inhibition of ICP1 (Fig. 1B-C), whereas DarTG exhibits potent anti-phage activity (22). The co-localization of Vc OLD and DarTG reflects the tendency of defense systems to cluster within genomic islands (42), potentially enabling them to work in concert to restrict phage infection (43, 44). When PDE^+^ *V. cholerae* are infected by ICP1, DarTG completely abolishes ICP1 replication and progeny production (22), likely precluding Vc OLD from functioning. However, when ICP1 isolates encode the DarT-inhibitor AdfB—a feature seen in clinical phage isolates following detection of the PDE (22)—Vc OLD would play a role in limiting phage propagation, providing additional protection to the bacterial population.

How Vc OLD is triggered to respond to phage infection remains an open question. Given that Vc OLD is not inherently toxic when expressed, we hypothesize that its nuclease activity requires activation during phage infection. Previously, it was unclear whether RecBCD is broadly required for the activation of Class 1 OLD family nucleases. Unlike P2 OLD, the only anti-phage Class 1 OLD family nuclease described previously, our data show that Vc OLD’s response to phage infection does not require host RecBCD (Fig. 4A-B). Furthermore, while P2 OLD inhibits lambda phage prior to genome replication (15), we find that Vc OLD-mediated inhibition of ICP1 occurs only after ICP1 initiates theta replication (Fig. 3). This suggests that perhaps Vc OLD activation is coupled to phage replication. Given that VC OLD restricts diverse vibriophages, we hypothesize that it is likely triggered by a global cellular perturbation during viral infection rather than a specific viral protein. A similar activation mechanism has been proposed for the Class 2 OLD family nuclease Bc GajA (*Bacillus cereus* GajA), which is thought to respond to perturbations of cellular nucleotide pools during phage infection (45, 46). Notably, Bc GajA nuclease activity is inhibited by high levels of ATP levels *in vitro*, a phenotype we did not see with Vc OLD (Fig. S5). As such, future studies are needed to discern the trigger(s) that mediate VC OLD’s response to phage infection.

Once active, VC OLD disrupts ICP1’s genome replication program (Fig. 3). Since Vc OLD can function as a nickase *in vitro* (Fig. 5), we hypothesize that it directly targets the replicating ICP1 genome *in vivo*, functioning similarly to Class 2 OLD-family nucleases (45, 46). Our results suggest that ICP1’s genome is degraded in the presence of Vc OLD (Fig. 3C). The mechanism by which nicking leads to degradation is unclear, but we speculate that indiscriminate Vc OLD-mediated nicking during infection may recruit additional host or phage-encoded nucleases. P2 OLD has been shown to have robust deoxyribonuclease activity and weak ribonuclease activity *in vitro* (21). Accordingly, following activation, P2 OLD could degrade both RNA and DNA in *E. coli,* resulting in cell death (16, 21). It remains to be elucidated whether Vc OLD directly acts on ICP1 DNA, and/or potentially targets RNA during infection. More broadly, the targets of OLD family nucleases are not yet fully understood, underscoring the need for further research to uncover their mechanisms and roles in phage defense.

Phages can escape defense systems by mutating the activator or target of the defense system or through gain-of-function mutations in inactive counter-defense proteins. In this study, we identified a phage-encoded direct inhibitor of the Class 1 OLD family nuclease, Vc OLD. Structural modeling of Vc OLD bound to Oad1 suggested that they likely interact at residues K58 and K166 of Vc OLD (Fig. 7A). Both residues fall within Vc OLD’s ATPase domain. Given this predicted interaction and the necessity of Vc OLD’s ATPase domain for its anti-phage activity *in vivo* (Fig. 4B), we hypothesize that Oad1 interferes with Vc OLD’s ATPase activity through direct binding. Specific inhibitors of ATPase domains as phage counter-defense mechanisms have yet to be documented (40), although a phage inhibitor that likely prevents the multimerization of a defense protein with an ATPase domain has been reported (47). Additional mechanisms are observed in other systems containing an ATPase domain-containing protein. For example, Gad1 (Gabija anti-defense 1) directly binds and encases the ATPase domain-containing Class 2 OLD family nuclease and downstream helicase (GajAB) complex, thereby blocking DNA recognition and cleavage (48).

All 67 ICP1 isolates in our collection encode a variant of Oad1 (37), but only four isolates from a discrete time period carry the functional counter-defense variant. This transient existence mirrors the fleeting presence of the PDE encoding Vc OLD. The pervasive presence of the non-functional variant of Oad1 raises questions about its potential role. While selection would favor maintenance of Oad1 if it served a critical function, we were able to delete it from ICP1’s genome without apparent deleterious effects on ICP1 fitness, at least under laboratory conditions. This suggests alternative reasons for the widespread maintenance of Oad1. Non-functional inhibitors that can be mutated into functional variants have been documented in other systems (22, 49, 50), supporting a model where such alleles provide a reservoir for rapid adaptation upon encountering a defense system. However, the rarity of the functional Oad1 variant and the failure of experimental evolution approaches to select ICP1 mutants that overcome Vc OLD suggest that the mutational pathway to a functional Oad1 variant may be constrained. Alternatively, the more common Oad1 variants could be functional against homologs of Vc OLD in under-surveyed environmental populations *V. cholerae*.

## Data Availability

The sequencing data generated for this work have been deposited in the Sequence Read Archive database under BioProject accession code PRJNA1186228.

## Author Contributions

Kishen M. Patel: Conceptualization, Investigation, Methodology, Formal analysis, Visualization, Validation, Writing—original draft. Kimberley D. Seed: Conceptualization, Supervision, Writing—review and editing, Funding acquisition.

## Acknowledgments

We thank past and present Seed lab members, especially Caroline M. Boyd and Reid T. Oshiro, for insightful discussion and feedback on this project. We also thank the helpful staff in the Vincent J. Coates Proteomics/Mass Spectrometry Laboratory Core Facility, RRID:SCR_025852.

