## Supplementary material for "A Class 1 OLD family nuclease encoded by *Vibrio cholerae* is countered by a vibriophage-encoded direct inhibitor": Table S1-S5, Figures S1-S8

**Table S1**: Bacteria and phage strains used in this study

*Numbers refer to the reference listed in the main text. References with roman numerals in parenthesis are additional references listed below the tables.

| Name in text | Strain | Description | Source* |
| --- | --- | --- | --- |
| Permissive PDE^-^ *V. cholerae* | KDS6 | *Vibrio cholerae* E7946, El Tor Ogawa, O1, streptomycin resistant | Levine et al., 1982 (I) |
| PDE^+^ *V. cholerae* | KMP268 | KDS6 containing the PDE integrated between VC0153 and VC0154 | Patel and Seed, 2024 (22) |
| PDE^+^ *V. cholerae* ∆*Vc old* | KMP298 | KMP268 with an in-frame frt cassette replacing *Vc old* | Patel and Seed, 2024 (22) |
| PDE^+^ *V. cholerae* ∆*Vc old* ∆*darT* | KMP393 | KMP298 with an in-frame frt cassette replacing *darT* | This study |
| PDE^-^ *V. cholerae* chromosomal expression *empty* | KMP594 | KDS6 with expression cassette (P_BAD_-riboswitchE) from Dalia *et al.*, 2020 (39)) integrated into *V. cholerae* lacZ locus with Kanamycin resistance cassette, expresses nothing (empty) | Patel and Seed, 2024 (22) |
| PDE^-^ *V. cholerae* chromosomal expression *Vc old* | KMP1177 | KMP594 with Vc OLD expressed from the expression cassette | This study |
| PDE^-^ *V. cholerae* chromosomal expression empty ∆*recBCD::specR* | KMP1207 | KMP594 with an in-frame spectomycin resistance marker flanked by two *frt* sites replacing *recBCD* | This study |
| PDE^-^ *V. cholerae* chromosomal expression *Vc old* ∆*recBCD::specR* | KMP1207 | KMP1177 with an in-frame spectomycin resistance marker flanked by two *frt* sites replacing *recBCD* | This study |
| PDE^-^ *V. cholerae* chromosomal expression *Vc old^K42A^* | KMP1209 | KMP594 with Vc OLD^K42A^ expressed from the expression cassette | This study |
| PDE^-^ *V. cholerae* chromosomal expression *Vc old^D313A^* | KMP1211 | KMP594 with Vc OLD^D313A^ expressed from the expression cassette | This study |
| PDE^-^ *V. cholerae* chromosomal expression *Vc old^D468A^* | KMP1213 | KMP594 with Vc OLD^D468A^ expressed from the expression cassette | This study |
| Empty Vector^+^ PDE^-^ *V. cholerae* chromosomal expression *empty* | KMP621 | KMP594 containing an empty vector | This study |
| p*gp106^2018^*^+^ PDE^-^ *V. cholerae* chromosomal expression *Vc old* | KMP1669 | KMP1177 containing a vector that can express Gp106^2018^ | This study |
| p*gp205^2018^*^+^ PDE^-^ *V. cholerae* chromosomal expression *Vc old* | KMP1185 | KMP1177 containing a vector that can express Gp205^2018^ | This study |
| PDE^-^ *V. cholerae* chromosomal expression *HA* | KMP1435 | KDS6 with expression cassette (P_BAD_-riboswitchE) from Dalia *et al.*, 2020 (39) integrated into *V. cholerae* lacZ locus with Kanamycin resistance cassette, expresses HA | This study |
| PDE^-^ *V. cholerae* chromosomal expression *Vc old-HA* | KMP1291 | KDS6 with expression cassette (P_BAD_-riboswitchE) from Dalia *et al.*, 2020 (39)) integrated into *V. cholerae* lacZ locus with Kanamycin resistance cassette, expresses Vc OLD with a C-terminal HA tag | This study |
| p*FLAG*^+^ PDE^-^ *V. cholerae* chromosomal expression *HA* | KMP1475 | KMP1435 containing a vector that expresses 3xFLAG | This study |
| p*FLAG*^+^ PDE^-^ *V. cholerae* chromosomal expression *Vc old-HA* | KMP1334 | KMP1291 containing a vector that expresses 3xFLAG | This study |
| p*gp205^2006^-FLAG*^+^ PDE^-^ *V. cholerae* chromosomal expression *Vc old-HA* | KMP1297 | KMP1291 containing a vector that expresses Gp205^2006^ with a C-terminal 3xFLAG tag | This study |
| p*gp205^2018^-FLAG*^+^ PDE^-^ *V. cholerae* chromosomal expression *Vc old-HA* | KMP1299 | KMP1291 containing a vector that expresses Gp205^2018^ with a C-terminal 3xFLAG tag | This study |
| p*FLAG*^+^ PDE^-^ *V. cholerae* chromosomal expression *empty* | KMP1332 | KMP594 containing a vector that expresses 3xFLAG | This study |
| p*FLAG*^+^ PDE^-^ *V. cholerae* chromosomal expression *Vc old* | KMP1187 | KMP1177 containing a vector that expresses 3xFLAG | This study |
| p*gp205^2018^-FLAG*^+^ PDE^-^ *V. cholerae* chromosomal expression *Vc old* | KMP1191 | KMP1177 containing a vector that expresses Gp205^2018^ with a C-terminal 3xFLAG tag | This study |
| p*gp205^2018_S136N^-FLAG*^+^ PDE^-^ *V. cholerae* chromosomal expression *Vc old* | KMP1659 | KMP1177 containing a vector that expresses Gp205^2018_S136N^ with a C-terminal 3xFLAG tag | This study |
| p*gp205^2018_K137D^-FLAG*^+^ PDE^-^ *V. cholerae* chromosomal expression *Vc old* | KMP1661 | KMP1177 containing a vector that expresses Gp205^2018_K137D^ with a C-terminal 3xFLAG tag | This study |
| p*gp205^2018_N144I^-FLAG*^+^ PDE^-^ *V. cholerae* chromosomal expression *Vc old* | KMP1663 | KMP1177 containing a vector that expresses Gp205^2018_N144I^ with a C-terminal 3xFLAG tag | This study |
| p*Vc old-HIS*^+^ Rosetta-gami B *E. coli* | KMP1427 | Rosetta-gami B *E. coli* containing a pETDuet vector that expresses Vc OLD with a C-terminal HIS tag | This study |
| ICP1^2006^ | KMPΦ36 | ICP1_2006_Dha_E (Accession Number: MH310934.1)  ∆*CRISPR* ∆*cas2-3* | McKitterick and Seed, 2018 (23) |
| ICP2 | ICP2 | Accession Number: HQ641345 | Seed et al., 2011 (36) |
| ICP3 | ICP3 | Accession Number: HQ641340 | Seed et al., 2011 (36) |
| ICP1^2006E^ ∆*gp205* | KMPΦ56 | KSΦ36 containing an in-frame deletion of *gp205,* host strain is KMP1127 | This study |
| ICP1_1992 | KMPΦ44 | ICP1_1992_Ind_M4 (Accession Number: MW794141) | LeGault et al., 2021 (41) |
| ICP1_2011 | KMPΦ1 | ICP1_2011 _A (Accession Number: MH310933) | Angermeyer et al., 2018 (II) |
| ICP1_2018 | KMPΦ47 | ICP1_2018_Mat _B (Accession Number: MW794183) | LeGault et al., 2021 (41) |
| ICP1_2018^1^ | PFS159 | ICP1_2018_Mat_159  (Accession Number: MW794177) | LeGault et al., 2021 (41) |
| ICP1_2018^2^ | PFS170 | ICP1_2018_Mat_170  (Accession Number: MW794182) | LeGault et al., 2021 (41) |
| ICP1_2019 | PFS242 | ICP1_2019_Mat _C (Accession Number: MW794190) | LeGault et al., 2021 (41) |
| ICP1_2019^1^ | PFS005 | ICP1_2019_Mat_005  (Accession Number: MW794188) | LeGault et al., 2021 (41) |

**Table S2**: Plasmids used in this study.

| Plasmids | Description and Identifier | Source |
| --- | --- | --- |
| Empty Vector | pMMB67EH engineered to contain an inducible riboswitch (E- induced by theophylline) downstream of a P*_tac_* promoter, KS1864 | Lab collection |
| p*gp106^2018^* | Empty vector engineered for the inducible expression of Gp106 from ICP1 2018, KMP1651 | This study |
| p*gp205^2018^* | Empty vector engineered for the inducible expression of Gp205 from ICP1 2018, KMP1077 | This study |
| p*FLAG* | Empty vector engineered for the inducible expression of 3x FLAG, CMB17 | Lab collection |
| p*gp205^2006^-FLAG* | pFLAG vector engineered for the inducible expression of Gp205^2006^ with a C-terminal 3xFLAG tag, KMP1139 | This study |
| p*gp205^2018^-FLAG*^+^ | pFLAG vector engineered for the inducible expression of Gp205^2018^ with a C-terminal 3xFLAG tag, KMP1143 | This study |
| p*gp205^2018_S136N^-FLAG* | pFLAG vector engineered for the inducible expression of Gp205^2018_S136N^ with a C-terminal 3xFLAG tag, KMP1642 | This study |
| p*gp205^2018_K137D^-FLAG* | pFLAG vector engineered for the inducible expression of Gp205^2018_K137D^ with a C-terminal 3xFLAG tag, KMP1644 | This study |
| p*gp205^2018_N144I^-FLAG* | pFLAG vector engineered for the inducible expression of Gp205^2018_N144I^ with a C-terminal 3xFLAG tag, KMP1646 | This study |
| p*Vc OLD-HIS* | pETDuet vector engineered for the inducible expression of Vc OLD with a C-terminal HIS tag, KMP1409 | This study |

**Table S3**: Percent Identity between varying gene products encoded by ICP1_2018 and the ICP1 isolates listed

|  | **ICP1 isolate** | | | | | | |
| --- | --- | --- | --- | --- | --- | --- | --- |
| **Gene Product** | 1992 | 2011 | 2006 | 2019 | 2018^1^ | 2018^2^ | 2019^1^ |
| Gp106 (protein ID: QVW06619.1) | 99 | 99 | 99 | 99 | 99 | 99 | 99 |
| Gp205 (protein ID: QVW06719.1) | 98 | 98 | 98 | 96 | 96 | 96 | 96 |
| Gp207 (protein ID: QVW06721.1) | 55 | 55 | 55 | 55 | 55 | 55 | 55 |

**Table S4**: List of proteins identified from Vc OLD-His purification

| Protein | Spectral Counts in Vc OLD-HIS Purification^1^ | Coverage (%) of Protein in Vc OLD-HIS Purification^2^ | Number of Peptides^3^ |
| --- | --- | --- | --- |
| Vc OLD-His | 280 | 36 | 36 |
| Elongation factor Tu 1 | 82 | 33 | 18 |
| Elongation factor Tu 2 | 82 | 33 | 18 |
| Chaperonin GroEL | 65 | 32 | 29 |
| Outer membrane porin F | 57 | 29 | 20 |
| Chaperone protein DnaK | 52 | 29 | 34 |
| Glyceraldehyde-3-phosphate dehydrogenase A | 39 | 24 | 16 |
| Elongation factor G | 33 | 19 | 15 |
| FKBP-type peptidyl-prolyl cis-trans isomerase SlyD | 28 | 15 | 2 |
| Alkyl hydroperoxide reductase C | 26 | 31 | 7 |

^1^Total number of peptides detected. Anything with a spectral count below 25 and coverage below 30 was removed from the table.

^2^Amount of protein sequence covered. Anything with sequence coverage below 30 and spectral count below 25 was removed from the table.

^3^Number of unique peptides detected.

**Table S5**: Average nucleotide identity between ICP1_2018 and the varying ICP1 isolates listed

|  | **ICP1 isolate** | | | |
| --- | --- | --- | --- | --- |
|  | 2019 | 2018^1^ | 2018^2^ | 2019^1^ |
| **Average nucleotide identity** | 99.965743 | 99.979743 | 99.978743 | 99.946143 |

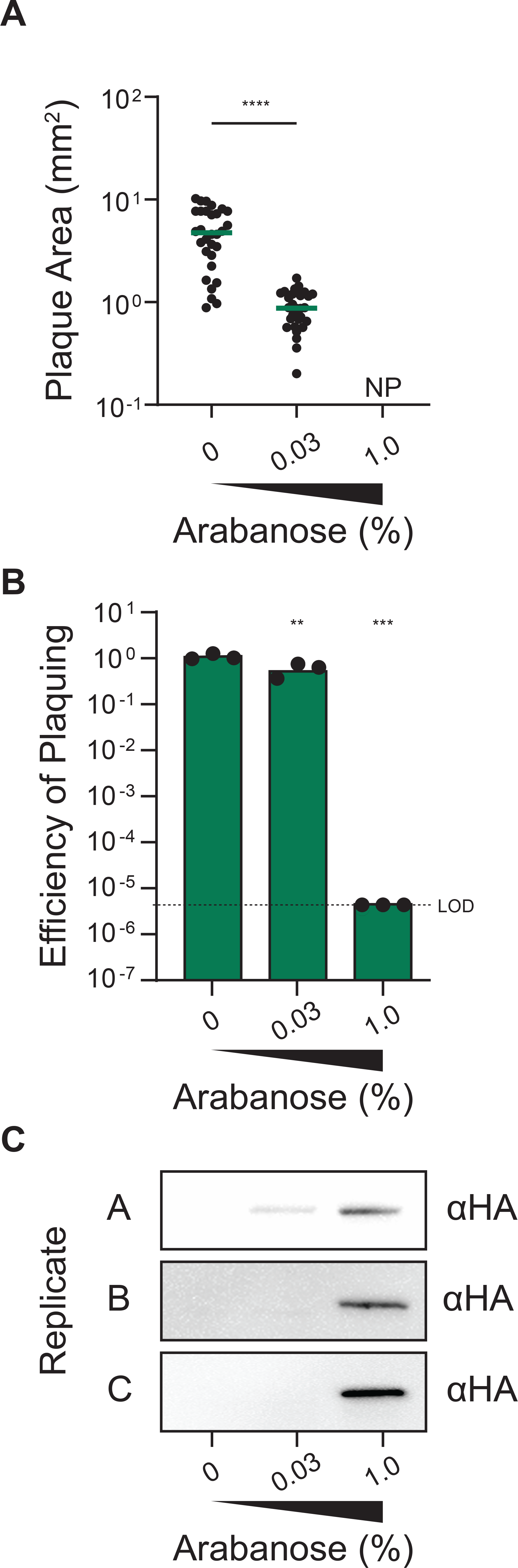

**Figure S1**: HA-tagged Vc OLD retains the capacity to inhibit ICP1 at higher induction conditions.

A) The plaque area (mm^2^) of ICP1 in *V. cholerae* expressing Vc OLD-HA at different percentages (w/v) of inducer (arabinose). NP represents no plaques. Each dot represents an individual plaque. Ten plaques were measured for each biological replicate for three biological replicates (n = 30). The green line indicates the mean of the plaque areas. Statistical analyses are results of unpaired *t*-test (*****P*<0.0001).
B) The efficiency of plaquing of ICP1 in *V. cholerae* expressing Vc OLD-HA at different percentages (w/v) of arabinose. Each dot represents an individual biological replicate. The bar indicates the mean of biological replicates. LOD represents the limit of detection. Statistical analyses are results of ANOVA followed by Dunnett’s T3 multiple comparison test using the 0% arabinose induction condition as the control condition (***P*=0.0085, ****P*=0.0002).
C) Western Blot against Vc OLD-HA before infection with ICP1 for plaque assay. Each biological replicate is shown on the left. Total protein from cell lysate was calculated and normalized to standardize total protein input (10µg) into each lane.

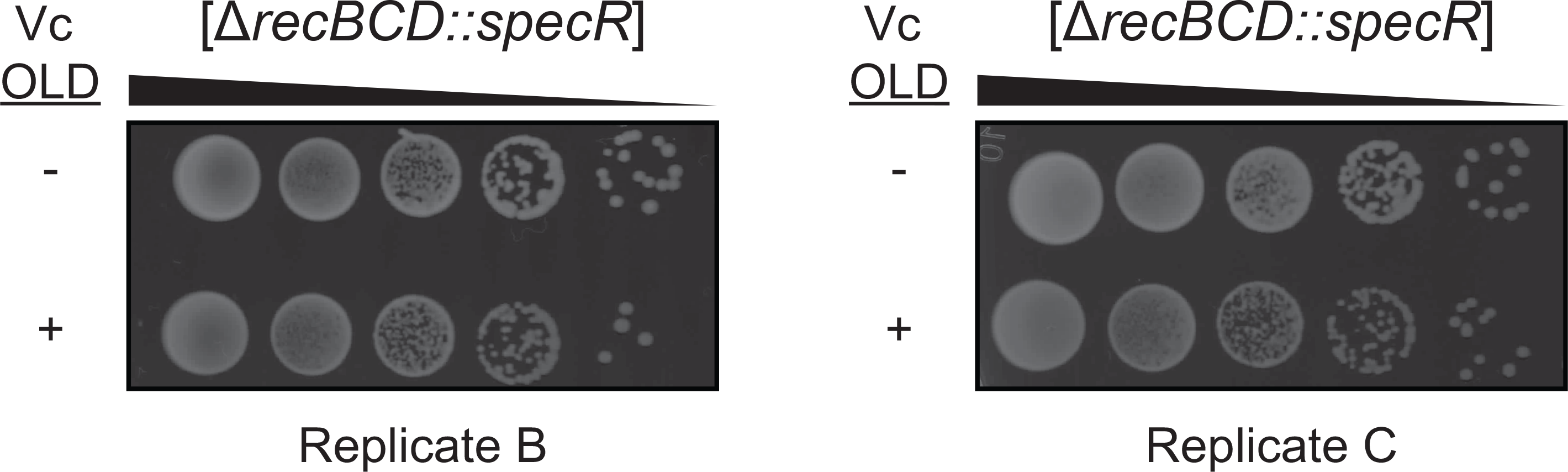

**Figure S2**: Biological replicates of toxicity assays of Vc OLD in RecBCD^-^ *V. cholerae* relative to an empty control.

Ten-fold serial dilutions of RecBCD^-^ *V. cholerae* expressing or not expressing Vc OLD. The black background is the agar plate, while the white spots are colonies. The replicate is written below the image.

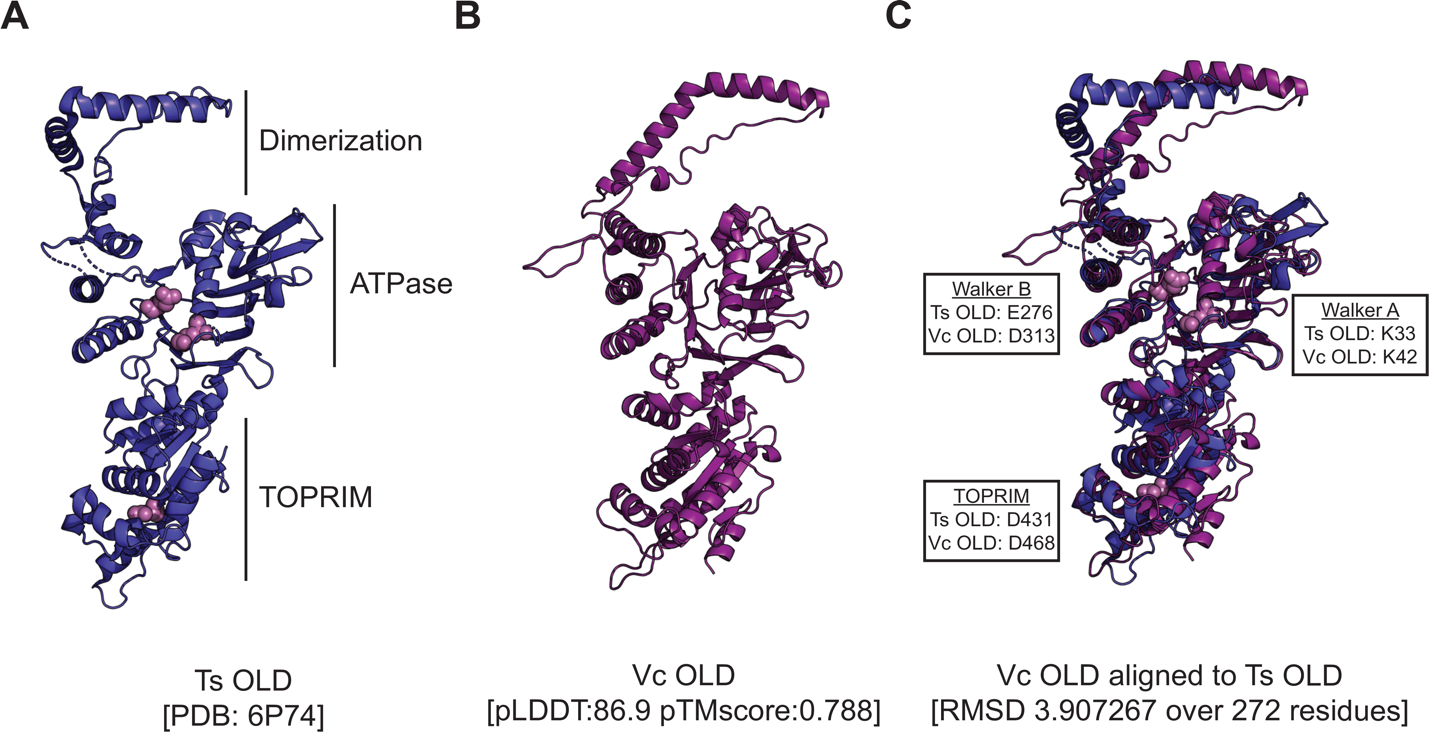

**Figure S3**: Ts OLD and the predicted structure of Vc OLD are similar.

A) The crystal structure of Ts OLD with domains labeled. In spheres and light purple are important residues for function *in vitro* (19).
B) The ColabFold predicted structure of Vc OLD from *V. cholerae*. pLDDT and pTM scores are written below the structures.
C) The superimposition of predicted structures of Vc OLD (dark purple) and Ts OLD (dark blue). The RMSD value is shown below the structure. Callouts show corresponding amino acids between Vc OLD and Ts OLD. When these residues are mutated for Ts OLD, they resulted in loss of function either by *in vitro* ATPase or Nuclease Assays (19).

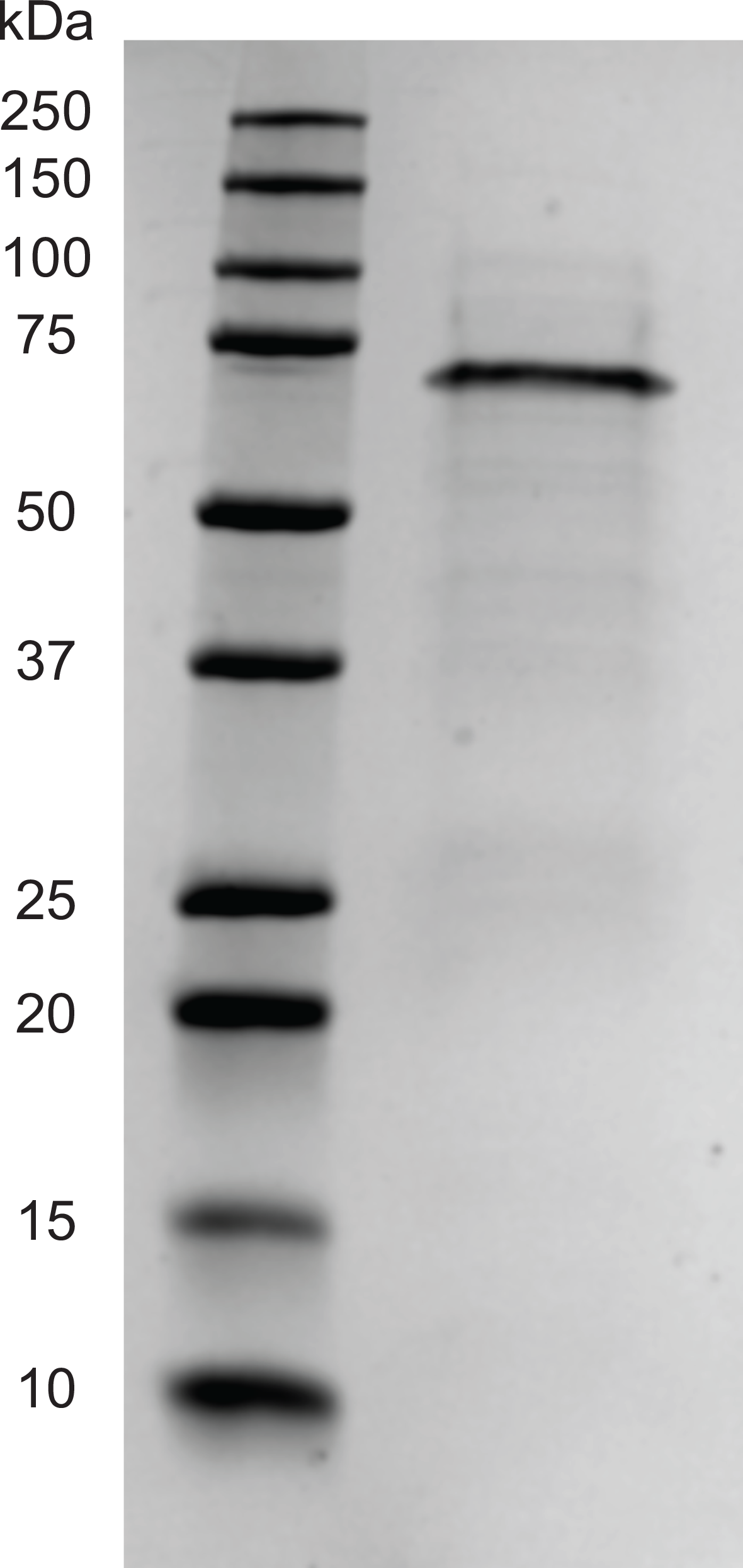

**Figure S4:** Coomassie-stained SDS-PAGE gel showing purified Vc OLD with a C-terminal HIS tag. Protein standard weights are listed on the left. Vc OLD with a C-terminal HIS tag has a molecular weight of ~66kDa.

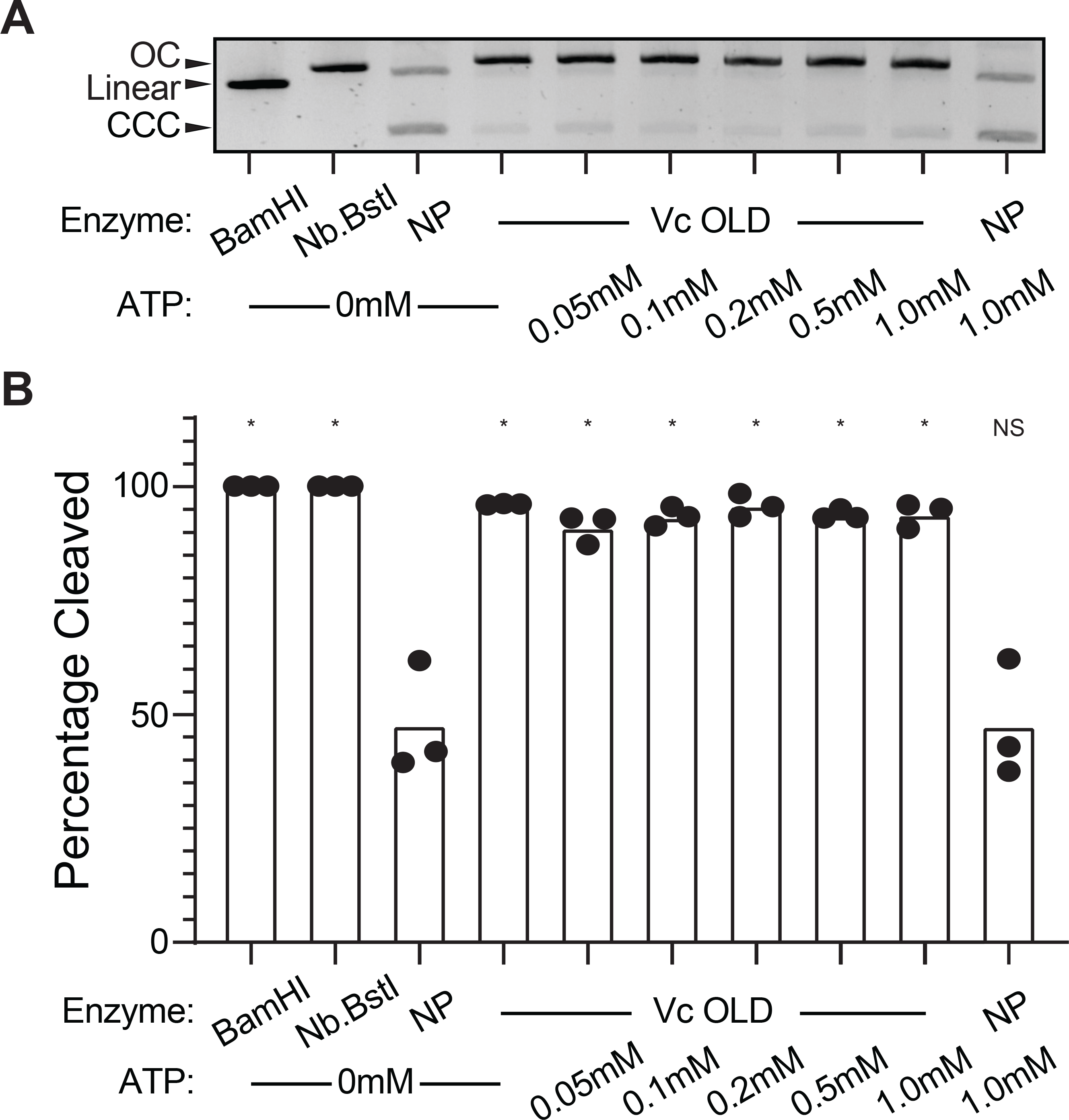

**Figure S5:** Vc OLD-mediated DNA nicking is unaffected by the addition of ATP.

A) *In vitro* cutting of a plasmid by 100nM of purified Vc OLD in the presence of varying amounts of ATP. Nicked or linearized plasmid substrate by restriction enzymes BamHI and Nb.BstI, respectively, are shown in the first two lanes. This image is representative of three replicates, which are quantified in panel B. NP stands for no protein.
B) Densitometry measurements calculating the percent of a plasmid cleaved by 100nM of purified Vc OLD in the presence of ATP at the concentrations listed. Each dot represents an independent replicate (n=3). The bar indicates the mean of replicates. Statistical analyses are the results of ANOVA followed by Dunnett’s T3 multiple comparison test using the no protein (NP) without ATP lane as the control condition (**P*<0.0001, NS = not significant).

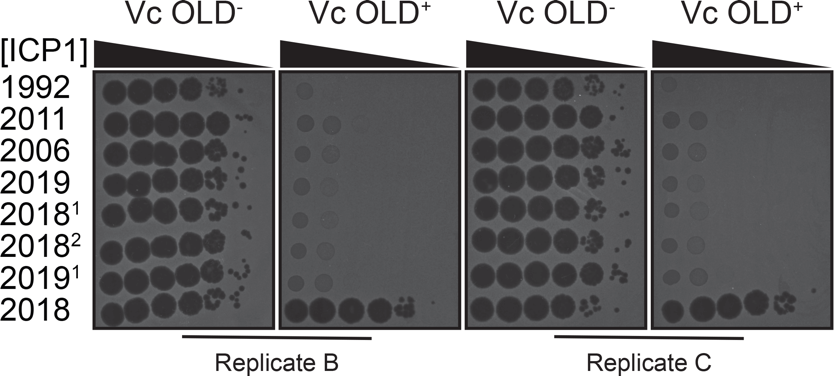

**Figure S6:** Biological replicates of ICP1 isolates indicated spotted on *V. cholerae* expressing or not expressing Vc OLD.

Ten-fold serial dilutions of the listed ICP1 isolate spotted on Vc OLD^+/-^ *V. cholerae* strains (right). Black zones of clearings are plaques. The opaque background is the *V. cholerae* lawn. The replicate is written below the plate.

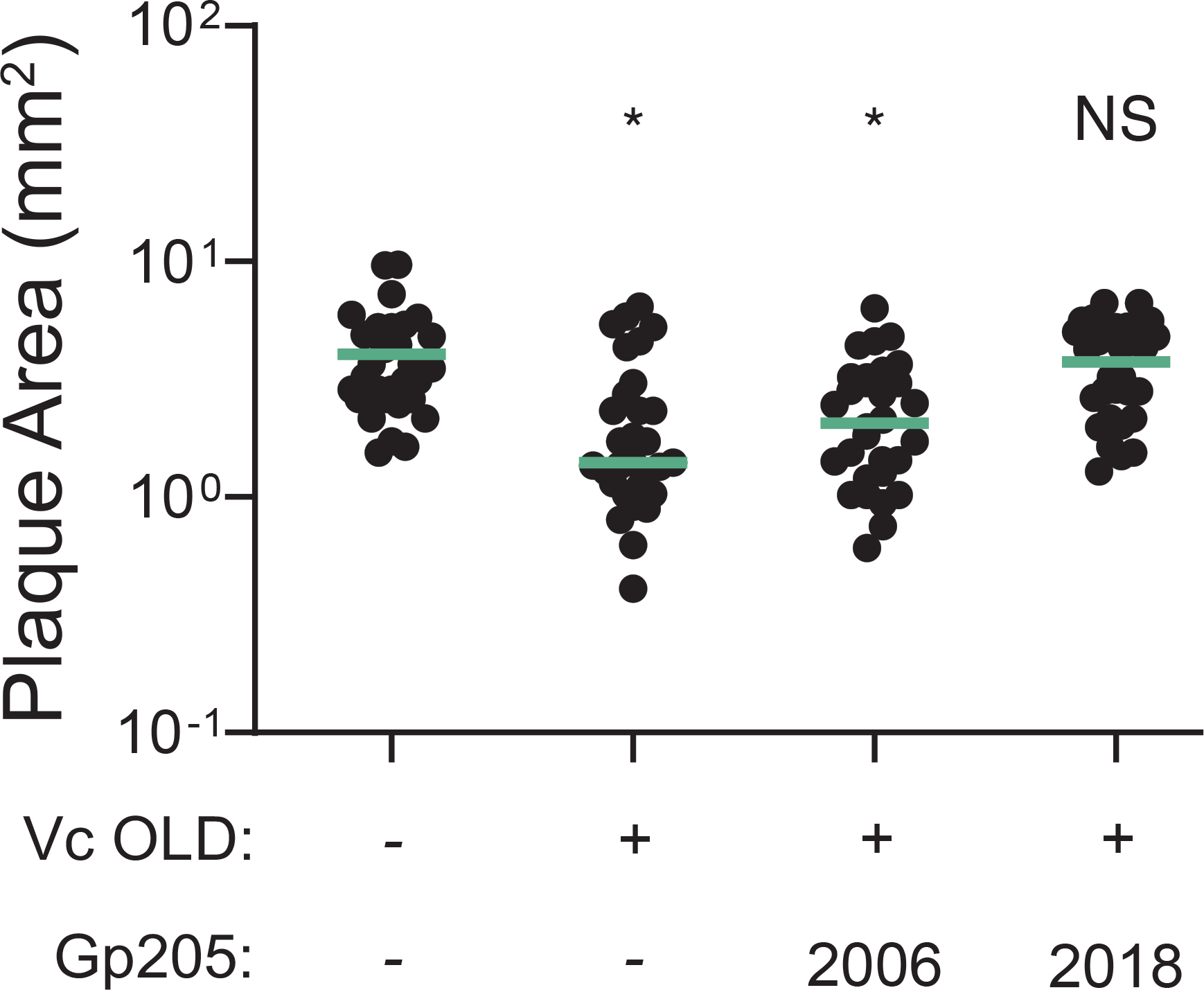

**Figure S7**: Gp205^2018^ restores ICP3 plaque size in the presence of Vc OLD.

The plaque area (mm^2^) of ICP3 plaques on Vc OLD^+/-^ expressing *V. cholerae* induced at 1% arabinose and an empty vector or a vector expressing Gp205^2006^ or Gp205^2018^. Each dot represents an individual plaque. Ten plaques were measured for each biological replicate for three biological replicates (n = 30). The green line indicates the mean of the plaque areas. Statistical analyses are the results of ANOVA followed by Dunnett’s T3 multiple comparison test using the empty vector containing Vc OLD^-^ *V. cholerae* strain as the control condition (*P<0.001, NS = not significant).

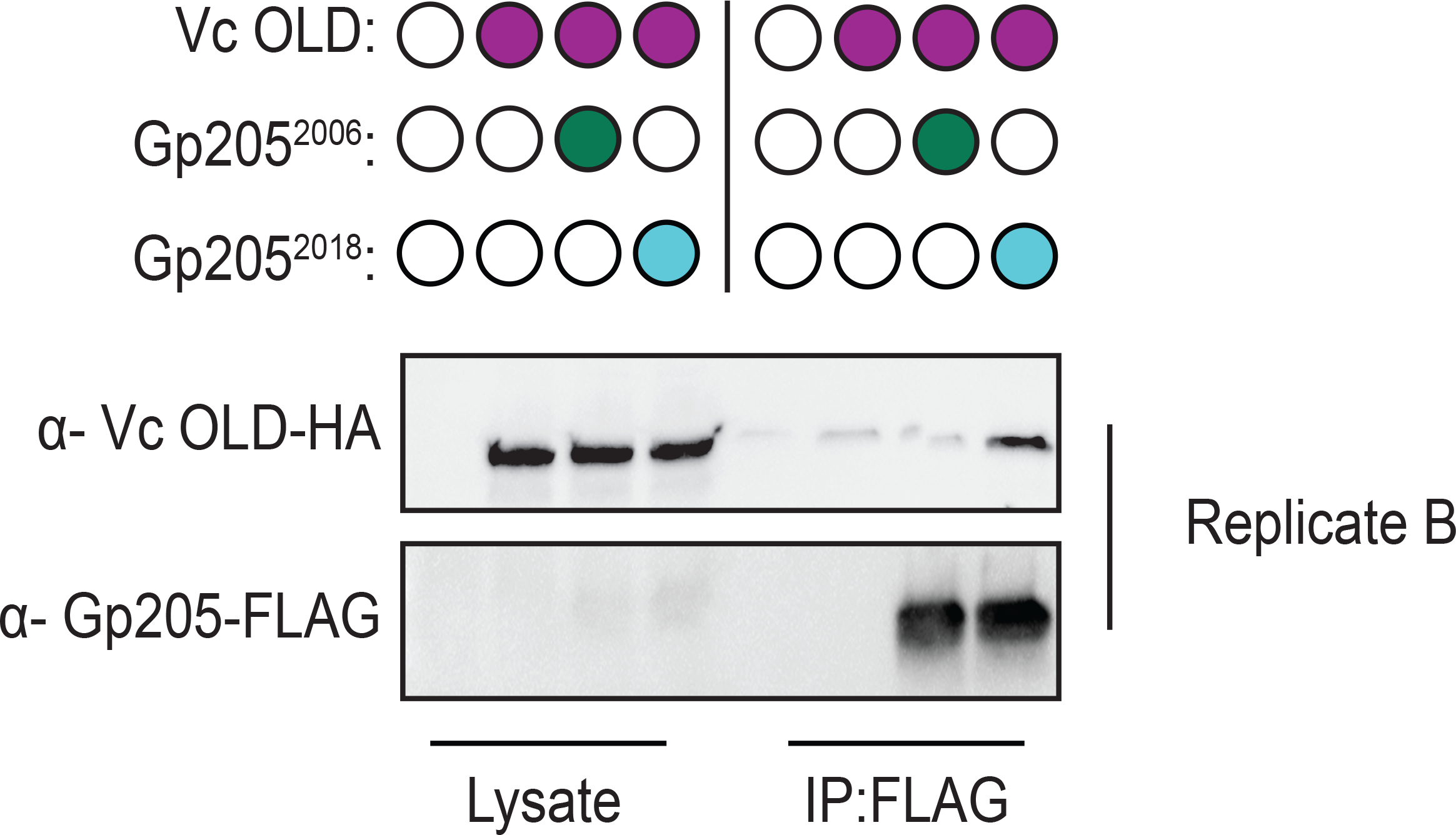

**Figure S8**: Western blot analysis of a biological replicate of the co-immunoprecipitation of Gp205-FLAG from cells expressing the proteins indicated (filled-in circle means being expressed). Lysates used as input for the immunoprecipitation (IP) were probed as controls for expression levels.
